# Efficient Bayesian Phylogenetics under the Infinite Sites Model

**DOI:** 10.1101/2025.11.14.688551

**Authors:** Ivan Specht, Julia A. Palacios

## Abstract

Bayesian phylogenetic inference from molecular sequences can provide key insights into the evolutionary history of populations. Existing tools, however, often scale poorly with sample size. We present `inPhynite,` a highly-efficient Bayesian phylogenetics algorithm for genomic datasets compatible with the infinite sites mutation model. A key advantage of this model is that likelihood calculation, which typically incurs a substantial computational cost, becomes trivial. We show that under the infinite sites assumption, it is possible to sample a coarse space of mutations and coalescences from which we may recover complete phylogenetic trees. We design an efficient Markov chain for this space together with effective population size trajectories, modeled as piecewise constant functions. Based on real and synthetic data, our method significantly outperforms competing methods, offering a speedup of over 225 times in statistical efficiency on large datasets without incurring any loss in accuracy. Finally, we demonstrate how `inPhynite` can help us understand the evolutionary history and past effective population sizes of human populations based on mitochondrial DNA.

**Summary:** Inferring the phylogenetic tree and evolutionary parameters from a sample of molecular sequences plays a key role in the study of how populations evolve over time. Existing inference algorithms face major computational challenges due to the large size of the phylogenetic tree space and high cost of phylogenetic likelihood evaluation. We show that under the infinite sites model of mutations, it is possible to overcome these limitations by instead conducting inference over an ordered sequence of genotypes that encodes the essential information in the tree. This approach achieves superior statistical efficiency compared to existing methods under a range of evolutionary conditions.

## 1 Introduction

The reconstruction of phylogenies and evolutionary parameters from molecular sequences is essential for a wide range of population-genetic and comparative analyses, such as identifying mutation events that occurred in the past, predicting the time of the most recent common ancestor of a set of taxa, and quantifying the change in population size for a species throughout history. Bayesian phylogenetics has become the state-of-the-art approach for reconstructing evolutionary histories because it quantifies uncertainty in both the tree topology and parameters, an advantage over maximum-likelihood methods that is especially important when genomic data are limited.

Numerous Bayesian methods for phylogenetic inference have been proposed, but many face major computational challenges that limit their utility. Most existing algorithms explore the posterior distribution via Markov chain Monte Carlo (MCMC), which iteratively makes minor alterations to the tree and evolutionary parameters. This approach has the drawback of sometimes getting “stuck” in local modes of the tree space and therefore can be slow to converge. This problem is only amplified under flexible models of the effective population size or for large input datasets, both of which result in a high-dimensional parameter space.

Considerable progress has been made recently in improving the practicality of Bayesian phylogenetics. A major advance came with `Delphy` (Varilly et al., 2025), which achieved statistical efficiency up to 1,000 times greater than that of `BEAST 2` (Bouckaert et al., 2014), a leading method, by augmenting the tree space with the mutations along each branch. At present, however, `Delphy` does not support population models besides constant and exponential growth, and the speedup was particularly marked for datasets with high overall sequence similarity. Other authors have explored alternatives to MCMC entirely, with a growing body of literature on Sequential Monte Carlo (SMC) and variational inference methods for tree sampling (Fourment and Darling, 2019; Zhang and Matsen, 2024). One such SMC-based approach proposes completely new trees at each iteration, building them up coalescence by coalescence, guided by regions of high probability density (Bouchard-Côté et al., 2012; Wang et al., 2015). Despite being parallelizable, SMC-based methods still suffer from a major inefficiency known as particle degeneracy, in which tree proposals are very often rejected due to insufficiently high probability (Wang and Wang, 2020; Milkey et al., 2025). Further refinements of SMC have also been proposed, such as SMC combined with annealed importance sampling (Wang et al., 2020), though preliminary results have also showed convergence challenges (Cappello et al., 2022). Variational inference methods can provide faster inferential results; however, these methods tend to underestimate uncertainty.

In this work, we design a highly efficient Bayesian phylogenetic inference method under an evolutionary model called the infinite sites model (or infinitely-many-sites model), which is relevant to a wide range of applications (Kimura, 1969; Watterson, 1975). A set of molecular sequences is compatible with the infinite sites model if no two mutations on the phylogeny ever occurred at the same site on the genome. Consequently, the model is particularly useful for analyzing datasets with high sequence similarity. It is also relevant to study large-scale molecular datasets: under this model, many phylogenies have likelihood 0, and so the infinite sites assumption can drastically shrink the state space of plausible trees (Palacios et al., 2022). When a dataset is compatible with the infinite sites model, it may be summarized as a perfect phylogeny, a graphical structure that captures the mutational history of the data and hence serves as a useful sufficient statistic for estimating the tree.

Phylogenetic inference under the infinite sites model has been a topic of study over the past several decades, and remains an active area of research. Griffiths and Tavaré (1994) formalized the process of coalescences and mutations under the infinite sites model, as well as proposed a Monte Carlo and an importance sampling method for estimating tree likelihoods. In Palacios et al. (2019) and Cappello et al. (2024), the authors reduced the latent tree space by modeling unlabeled ranked tree shapes instead of labeled coalescent trees; however, this comes with a higher likelihood cost as its calculation needs to consider all possible allocations of mutations to branches. More recently, Holbolth et al. showed that there exists a matrix-analytical exact formula for sampling infinite sites trees, though it is tractable to compute only for small total numbers of mutations due to the combinatorially-large size of the space of trees and mutations (Hobolth et al., 2025).

We advance the state of the art in infinite sites Bayesian phylogenetics in two key ways. First, we leverage the infinite sites assumption to model not only the latent phylogenetic structure, but rather the phylogeny annotated with mutations in such a way that it is compatible with the observed data in the form of a perfect phylogeny. We then show that it is possible to compress these annotated trees into objects called *evolutionary paths*, which store the ordered sequence of genotypes in which each coalescence and mutation occurs. For inference, we target the posterior distribution of evolutionary paths and population sizes, which offers the key advantages of (1) the likelihood calculation becomes trivial, and (2) an evolutionary path sample from the posterior may be converted to a phylogenetic tree sample by a simple random generation step. This approach achieves significant computational gains against other methods because the space of evolutionary paths can be sampled far more efficiently than the entire space of phylogenies. Finally, we show that by working with this highly compressed tree resolution, Bayesian non-parametric estimation of the effective population size can be made efficient by assuming the change points agree with coalescent times, which is the typical assumption in “Skyline” methods (Drummond et al., 2005; Minin et al., 2008; Ho and Shapiro, 2011). This assumption allows us to integrate out branch length information and design a Markov chain of evolutionary paths and effective population sizes, independent of coalescent times.

We combine our tree and effective population size samplers to arrive at our method, `inPhynite`. We demonstrate that `inPhynite` significantly outperforms `BEAST 2` in terms of statistical efficiency (effective sample size per unit runtime) based on real and synthetic datasets—without incurring any loss in accuracy. We show the speedup to be particularly drastic on large datasets with many mutations, indicating superior scalability for `inPhynite` as compared to `BEAST 2`. By applying `inPhynite` to mitochondrial DNA datasets from the 1,000 Genomes Project, we demonstrate how our method can be used to infer the effective population size of human groups over time.

## 2 Constructing Evolutionary Paths

### Overview

We first review the standard (Kingman) coalescent on timed, labeled, binary trees. We then introduce the infinite sites model, assuming a Poisson process for the mutations along each tree branch. This model assumption gives rise to a lossless representation of the data as a perfect phylogeny and the closely related structure of an augmented perfect phylogeny, which explicitly accounts for all observed and unobserved genotypes in the evolutionary history of the sample. This representation is particularly useful as it delimits the space of possible genotypes that may occur on any inferred phylogeny. Next, we provide an alternative characterization of the joint probability of the coalescent tree and mutational history. This characterization allows us to lump together trees based on whether they share the same order of coalescent and mutation events (i.e. the same embedded jump chain), and based on whether said events involve the same genotypes. Finally, conditioning on the observed augmented perfect phylogeny, we arrive at a resolution we call an *evolutionary path* consisting of only the ordered sequence of genotypes in which each coalescence/mutation occurs. See Figure 1 for an overview of tree resolutions and the construction of evolutionary paths. Despite its coarseness, posterior samples at this resolution (jointly with the effective population size) are all that are needed to obtain posterior samples of timed phylogenetic trees. This conversion can be achieved via a post-processing step that involves sampling the branch lengths and topology from simple, analytic probability distributions.

**Figure 1:**
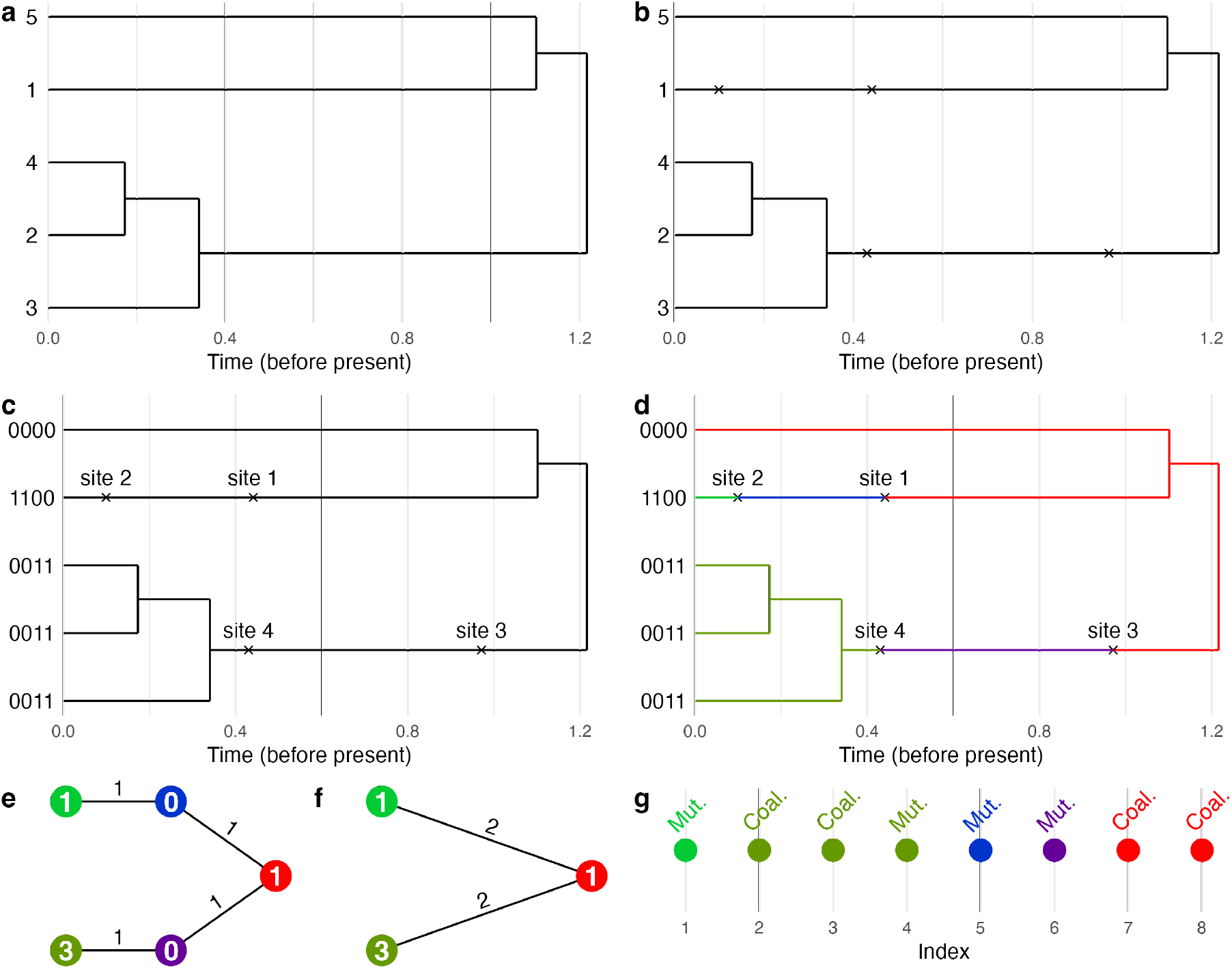
The construction of the evolutionary path. (a) We start with a timed, leaf-labeled phylogenetic tree. (b) We annotate each branch of the tree with mutations, sampled according to a Poisson process. (c) We assign to each mutation a different site on the genome that it affects, uniformly at random. (d) These mutations allow us to partition the tree into different connected regions (shown in different colors), each of which has its own unique genotype. (e) From a partitioned tree, we may extract the adjacency graph of the colored regions, which is called an *augmented perfect phylogeny*. The node color corresponds to the genotype of a region, and the node weight (number on the node) represents the number of sampled leaves of that genotype. (f) An augmented perfect phylogeny may be transformed into a *perfect phylogeny* by collapsing internal nodes with degree 2 and weight 0. The edge weights represent the number of mutations separating a pair of genotypes. (g) An *evolutionary path* is a sequence whose *i*th element is the genotype involved in the *i*th event (coalescence or mutation).

### 2.1 Timed, Labeled Trees

The first class of phylogenetic trees we consider are timed, labeled trees. Labeled trees, also known as Kingman trees, are trees in which the *n* leaves are each assigned a unique identifier among the integers 1, 2, …, *n*. The word *timed* refers to the assignment of each coalescence (internal node) a time in ℝ_+_, with time measured in units since the present. A visualization of a timed, labeled tree is shown in Figure 1a.

The probability density assigned to a timed, labeled tree under Kingman’s coalescent with variable effective population size is expressed as a product of conditional densities, each representing the density of the waiting time until the next coalescent event. Let 0 = *t*_0_ *< t*_1_ *< t*_2_ *<* …, *t*_*n−*1_ be the coalescent times and let *N* : ℝ_+_ → ℝ_+_ be the effective population size, a time-dependent quantity representing the number of reproducing individuals in the population scaled by the duration of a generation. Let *k*_*i*_ := *n −*(*i*)+ 1, the *number of extant lineages* right before the *i*th coalescent event, for *i* ≥ 1, and let 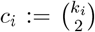. The probability density for a tree *x* with a particular topology and coalescent times (*t*_0_, …, *t*_*n−*1_) is then given by

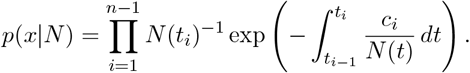

In this paper, we are particularly interested in the special case in which *N* (*t*) only changes at coalescent times, and is otherwise constant. This assumption, which may be viewed as a piecewise-constant approximation to general *N* (*t*), facilitates statistical inference while still allowing us to model flexible, non-parametric effective population size trajectories. It is a core idea behind two standard non-parametric models implemented in `BEAST 2`, “Skyline” and “Skyride” (Minin et al., 2008; Drummond et al., 2005). With *N* (*t*) constant on each intercoalescent interval, let *N*_*i*_ denote the value of *N* on the interval (*t*_*i−*1_, *t*_*i*_]. The probability density then becomes

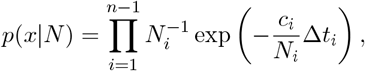

where Δ*t*_*i*_ := *t*_*i*_ *− t*_*i−*1_.

### 2.2 Mutations and Sequence Data

The previous section establishes a prior distribution over phylogenies; we now show how a Poisson process of mutations may be superimposed on the tree to generate molecular sequences at the leaves. Given a timed, labeled tree, we model the number of mutations *m*_*i*_ along a branch of the tree of length *ℓ*_*i*_ independently according to a Poisson distribution with rate parameter *µℓ*_*i*_, where *µ* is the mutation rate, treated as fixed and known (we show in Section 2.5 that leaving *µ* as an unknown parameter leads to identifiability issues). Conditional on the number of mutations, the mutation times are uniformly distributed along each branch (see Figure 1b). Mutation times are not explicitly needed to obtain the molecular sequences at the leaves; however, we do need to specify at which site along the genome each mutation event occurs. Let

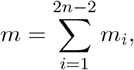

denote the total number of mutations, which equals the total number of polymorphic sites of the samples. The sum ranges from *i* = 1 to 2*n* − 2 because a tree with *n* leaves has 2*n* − 2 total branches. We select a uniformly random bijective map *ϕ* that assigns to each of the *m* mutations on the tree a site *j* ∈ {1, 2, …, *m*} . The probability mass function of *ϕ*, conditional on the number of mutations on each branch, is hence given by

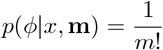

where **m** = (*m*_1_, …, *m*_2*n−*2_). The joint mass of *ϕ* and **m** given *x* is therefore

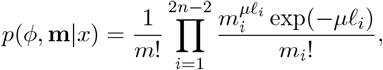

with the terms inside the product being each of the Pois(*µℓ*_*i*_) probability masses. We then define the sample *s*_*i*_ at leaf *i* to be a binary sequence of length *m*, whose *j*th element 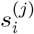 equals 1 if mutation *j* is in the ancestral history linking leaf *i* back to the root, and 0, otherwise. Figure 1c shows an example of a phylogeny annotated with mutations, the mapping of mutations to sites, and the corresponding tip sequences.

The previous paragraphs explain how to generate molecular sequences (at polymorphic sites) under the infinite sites model, given a tree *x*. We now discuss how to calculate the likelihood of a tree annotated with mutations, given data in the form of molecular sequences observed at the tips. First, we filter the sequence dataset *S* = (*s*_1_, …, *s*_*n*_) to polymorphic sites, as our model quantifies the likelihood associated with all polymorphic (as opposed to conserved) sites on the genome. We next check that the sequences are indeed compatible with the infinite sites model (see, e.g., Griffiths and Tavaré (1994) and Gusfield (1991)). If they are not, we return a likelihood of zero (in practice, we may apply a masking procedure to the sequences prior to inference to avoid this scenario). The probability mass function of the data *S* given *x*, **m**, and *ϕ* is given by

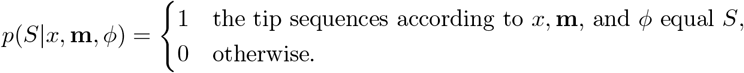

We emphasize that to calculate the likelihood, it is enough to know which mutations occurred on each branch, and not the time at which the mutation events occurred.

We note one final subtlety associated with this likelihood function: the sequences produced by our generative model are expressed as changes relative to the sequence at the root of the tree. If the genotype of the root is specified by the user prior to inference, there is no ambiguity in our likelihood. If, however, the root state is left unspecified, the definition of *p*(*S*|*x*, **m**, *ϕ*) is in fact the conditional likelihood of (*x*, **m**, *ϕ*) given the root state. Assuming a uniform prior over root states compatible with the infinite sites model, the likelihood remains valid as stated, up to a constant of proportionality.

### 2.3 Perfect Phylogenies as Sufficient Statistics

While Section 2.2 establishes a perfectly valid model for trees and molecular sequences, repeatedly checking whether leaf sequences according to (*x*, **m**, *ϕ*) agree with *S* is expensive, and it is unclear how best to explore the space of (*x*, **m**, *ϕ*) with nonzero likelihood. To help make statistical inference more tractable, we construct two convenient representations of molecular data generated under the infinite sites model known as a *perfect phylogeny* and an *augmented perfect phylogeny*. We define the latter first, from which it is straightforward to obtain the former. An augmented perfect phylogeny is an undirected tree in which each node represents a unique genotype that appeared at some point in the evolutionary past, and an edge connects two nodes (genotypes) if one evolved from the other by way of a single mutation event. Note, crucially, that some of the nodes on an augmented perfect phylogeny may represent *unknown* genotypes, and merely reflect the fact that *some* genotype existed at one point in the evolutionary past (we expand on this point momentarily).

Each node in an augmented perfect phylogeny is assigned a weight equal to the number of observed sequences whose genotype corresponds to that node. The leaf nodes of the augmented perfect phylogeny necessarily have positive weight, as they correspond to one or more observed sequences, while the other nodes need only have nonnegative weight. Additionally, if the user wishes to fix the ancestral genotype of the sample of sequences (in case this is known), the node corresponding to said genotype is marked as the “root” node of the augmented perfect phylogeny. A *perfect phylogeny*, then, is the same as an augmented perfect phylogeny, except that internal nodes with degree 2 and weight 0 on the augmented perfect phylogeny are collapsed (see procedure below for details and Figure 1e–f). Unlike in an augmented perfect phylogeny, every node in a perfect phylogeny corresponds to a unique, known genotype (though not necessarily an observed one).

Suppose we are given a tree *x*, the number of mutations **m** on each branch, and a bijection *ϕ* assigning to each mutation a position *j* ∈ {1, …, *m*} among the *m* polymorphic sites on the genome. In this context, it is straightforward to obtain the corresponding augmented perfect phylogeny and standard perfect phylogeny. A simple algorithm for doing so is as follows:

1. Create a node for each unique genotype on the tree, including those that do not have any representative leaves. In Figure 1d, these unique genotypes are shown by the five colors on the tree. Set the weight of each node to be the number of leaves of the corresponding genotype. Create an edge between two nodes if there exists a mutation between their corresponding genotypes on the tree. Set the weight of each edge to be 1. This gives us our augmented perfect phylogeny (Figure 1e).
2. Identify a node that (a) has zero representative leaves, (b) has exactly two edges connected to it, and (c) is not the root node, if the user wishes to conduct inference with a specified root genotype. Remove the node and wire together its two neighbors. The weight of the newly created edge is the sum of the weights of the two deleted edges.
3. Repeat step 2 until no longer possible. Once done, we obtain the perfect phylogeny (Figure 1f).

The above gives a procedure for obtaining the (augmented) perfect phylogeny from a known tree *x* with mutation counts **m** and mutation-to-site mapping *ϕ*. We now turn to the question of constructing an (augmented) perfect phylogeny from a dataset *S* of sequences alone. Remarkably, any dataset of genome sequences compatible with the infinite sites model implies a unique underlying perfect phylogeny (Palacios et al., 2019). Algorithms for converting a set of genome sequences to its perfect phylogeny have been proposed by, e.g., Griffiths and Tavare (1994) and Gusfield (1991) and are not discussed here. We note that in our construction of perfect phylogenies, we collapse leaf edges with weight 0 (in contrast to, e.g., Palacios et al. (2019)).

By expanding each edge of weight *w* ≥ 2 on said perfect phylogeny with *w* consecutive edges of weight 1, interspersed with nodes of weight 0, we obtain a unique augmented perfect phylogeny from the data (see conversion of Figure 1f to Figure 1e as an example). We stress that each new node created via this process does *not* correspond to any one particular genome sequence, but rather reflects the fact that *some* intermediate genotype(s) must have existed to separate two nodes connected by an edge of weight *w* ≥ 2 on the perfect phylogeny. For example, in Figure 1d, it is not possible to tell from the data whether the blue genotype in panel (b) represents the genome sequence `1000` or `0100`. However, it is known from the data that the light green genotype evolved from the red genotype (or vice versa) by way of two mutations, occurring at sites 1 and 2 (with their order unknown). We will not need augmented perfect phylogenies for the remainder of this section, but they will prove crucial for designing an inference algorithm later on.

We claim that a perfect phylogeny (and hence also an augmented perfect phylogeny) is a sufficient statistic for (*x*, **m**). To prove this, we show that there exist functions *g, h* such that

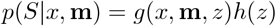

where *z* is the perfect phylogeny obtained from observed data *S* and *h* is a function only of *z*. To see this, we expand the left-hand side by conditioning on *ϕ*:

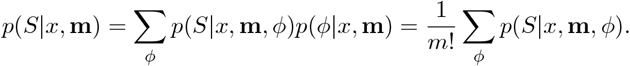

where the sum ranges over all bijective mappings from the set of mutations on the tree to 1, …, *m* . Suppose there exists some *ϕ* such that *p*(*S*|*x*, **m**, *ϕ*) = 1. This happens exactly when the perfect phylogeny obtained from (*x*, **m**) is graph-isomorphic to the perfect phylogeny obtained from *S*, where the isomorphism preserves node weights and (optional) root node. One can see that for any branch *i* on the tree *x*, permuting the site labels under *ϕ* of the *m*_*i*_ mutations along that branch has no effect on the likelihood *p*(*S*|*x*, **m**, *ϕ*), but swapping site labels between branches will result in a likelihood of 0. Hence, there are exactly

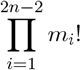

terms in the summation for which *p*(*S*|*x*, **m**, *ϕ*) = 1; for all others, *p*(*S*|*x*, **m**, *ϕ*) = 0. Therefore, so long as there exists some *ϕ* with *p*(*S*|*x*, **m**, *ϕ*) = 1, we have

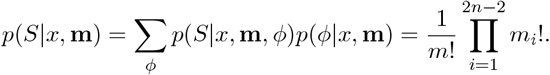

Note that the above is in fact a function of *z*, as the *m*_*i*_’s are in bijection with the edge weights of *z* and *m* is the sum of the *m*_*i*_’s. Hence, we take

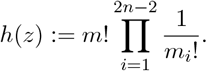

Letting

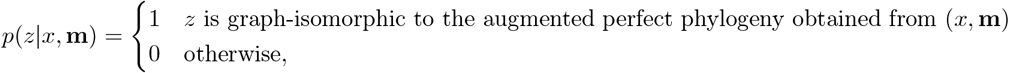

we obtain

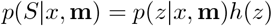

and therefore *z* is sufficient for (*x*, **m**). Moving forward, we may thus treat our data as a perfect phylogeny *z* instead of sequences *S*, and work with the likelihood *p*(*z*|*x*, **m**) instead of *p*(*S*|*x*, **m**).

### 2.4 Timed, Labeled, Mutation-Annotated Trees

The previous sections show that computing the likelihood of a mutation-annotated tree reduces to a matter of checking whether it is compatible with the observed perfect phylogeny. With this in mind, it is useful to consider the tree and mutations together as one joint process. Define a timed, labeled, mutation-annotated tree (TLMAT) to be the same as a timed, labeled tree, except that the time of each mutation event is also represented explicitly. An example of a TLMAT is shown in Figure 1b.

To define a prior probability density on the space of TLMATs, we follow the work of Griffiths and Tavaré (1994). We start with the continuous-time Markov chain of coalescent and mutation events going backwards in time from the present, from which we then derive the probability of the TLMAT. Let 0 = *t*_0_ *< t*_1_ *<* · · · *< t*_*m*+*n−*1_ represent the times of all coalescences and mutations, where *m* is the total number of mutation events on the tree. Let *χ*_*i*_ = 1 if time *t*_*i*_ corresponds to a coalescence and 0 if time *t*_*i*_ corresponds to a mutation. As before, let Δ*t*_*i*_ = *t*_*i*_ *™ t*_*i−*1_ for *i ≥* 1. Since the mutations along each branch follow a Poisson process with rate *µ*, the waiting time to the next mutation event when there are *k* extant branches is exponentially distributed with rate *kµ*. Moreover, from our tree prior, we know that the waiting time to the next coalescent event when there are *k* extant branches is exponentially distributed with rate 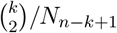. Hence, the transition density of the chain (Δ*t*_*i*_, *χ*_*i*_) is given by

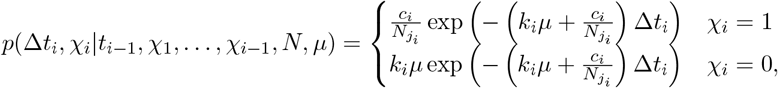

where *k*_*i*_ denotes the number of extant lineages immediately before time *t*_*i*_, now equal to

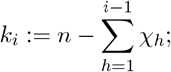

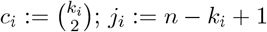; and *p*, with slight abuse of notation, denotes both the continuous (over Δ*t*_*i*_) probability density and discrete (over *χ*_*i*_) probability mass. Given the chain 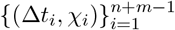, a TLMAT *w* is then generated by sequentially sampling the single extant lineage (in the case of a mutation event) or the pair of extant lineages (in the case of a coalescence) involved at each event, uniformly at random. The resulting density of a TLMAT *w* is given by

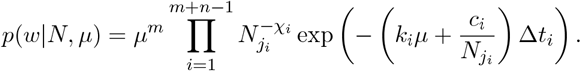

### 2.5 Ranked, Labeled, Mutation-Annotated Trees

Recall that a TLMAT records the events and times in which coalescences and mutations occur on a tree. Conveniently, under the assumption that *N* (*t*) is constant on each intercoalescent interval, it is possible to lump together all TLMATs that are the same up to the waiting times Δ*t*_*i*_ between subsequent coalescences and mutations, and still obtain an analytic form for the probability distribution of the resulting equivalence classes of TLMATs.

To formalize this idea, we define a *ranked, labeled mutation-annotated tree* (RLMAT) to be an equivalence class of TLMATs that differ only in the values of the Δ*t*_*i*_’s. The word *ranked* here refers to the fact that in a RLMAT, we store the order but not the exact time of each coalescence or mutation. We write V_*m,n*_ to denote the space of all RLMATs with *n* leaves and *m* mutations. We express the probability density of a RLMAT *v* by integrating over all such *w* that are in the same equivalence class:

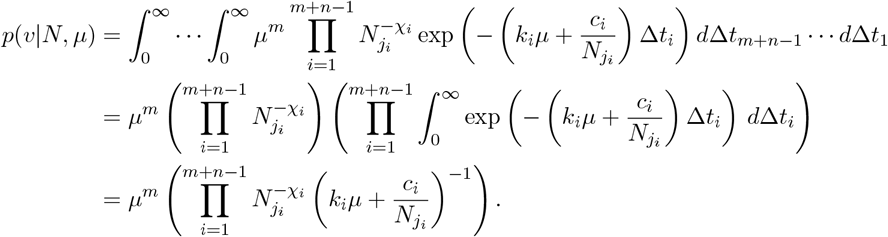

We remark here that *µ* and *N* are not identifiable from one another. To see this, we multiply the right-hand side by 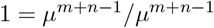, and invoke the fact that *χ*_*i*_ = 1 for exactly *n* − 1 values of *i*, to obtain

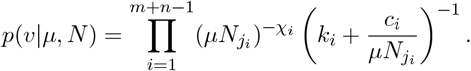

From this, we see that *p*(*v*|*µ, N*) can be expressed as a function of the vector *µN* . Moving forward, we may assume without loss of generality that *µ* = 1 deterministically, and we will infer *N* relative to *µ*. Hence, we may write the probability distribution over RLMATs *v* as

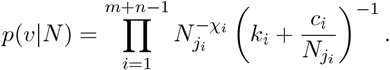

### 2.6 Evolutionary Paths

So far, we have defined a prior density on RLMATs in the absence of any data. Assuming the infinite sites mutation model and conditioning on the data *z* in the form of an (augmented) perfect phylogeny, it is further possible to lump together RLMATs into a still coarser tree resolution that permits highly efficient sampling. This coarser tree resolution, which we call an *evolutionary path*, encodes the ordered sequence of genotypes in which each coalescence or mutation occurs. In contrast to an RLMAT, an evolutionary path does *not* record the whole labeled tree topology, i.e. for each coalescence, it does not record which specific pair of lineages coalesce. Figure 1g shows the evolutionary path corresponding to the example tree used throughout this section, with each genotype represented by a different color.

To extend the probability measure *p* from RLMATs to evolutionary paths, let 𝕌_*z*_ denote the space of all evolutionary paths compatible with *z*. We can represent an element *u* ∈ 𝕌_*z*_ as

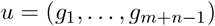

where *g*_*i*_ represents the genotype on the augmented perfect phylogeny in which the *i*th coalescence or mutation occurs, in order from earliest (closest to leaves) to latest (closest to root). A mathematically rigorous method for obtaining the types (coalescence or mutation) from an evolutionary path and the perfect phylogeny is provided in Appendix A, Corollary 1.

To lump together RLMATs that share the same evolutionary path, we define an equivalence relation ∼ _*u*_ on 𝕍_*m,n*_ where *v* ∼ _*u*_ *v*^*′*^ if—conditional on the perfect phylogeny *z*—the genotype in which the *i*th coalescence or mutation occurs in RLMAT *v* equals the genotype in which the *i*th coalescence or mutation occurs in RLMAT *v*^*′*^. The cumulative probability assigned to all *v* in an equivalence class *u* ∈ 𝕌_*z*_ = 𝕍_*z*_*/* ∼ _*u*_ is obtained by counting the number of possible pairs of extant lineages that may coalesce in a given genotype:

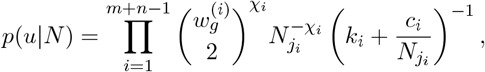

where 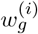 is the number of extant lineages of genotype *g*_*i*_ when the *i*th coalescence or mutation occurs, and it is assumed above that *u* is compatible with the data *z* in the form of a perfect phylogeny. Note that the indicators *χ*_1_, …, *χ*_*m*+*n−*1_ are not included in the definition of an evolutionary path, as they can be reconstructed deterministically from *z* and *u*: *χ*_*i*_ is a coalescence if and only if there exists *j > i* such that *g*_*j*_ = *g*_*i*_ (see Appendix A for justification). We develop a computationally efficient procedure for checking whether an evolutionary path is indeed compatible with a perfect phylogeny in Appendix A. This space 𝕌_*z*_—the sequence of genotypes in which each coalescence or mutation occurs, independently of the times—is what we sample explicitly with `inPhynite`.

In Section 3, we develop a procedure to sample evolutionary paths and population sizes from the posterior distribution. Then, given a posterior evolutionary path, effective population size trajectory, and perfect phylogeny, we can sample from the posterior distribution over timed, labeled trees as follows:

1. Start with an empty tree consisting of just the leaves. For each genotype *g* on the perfect phylogeny, assign 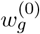 of the leaves to be of genotype *g*. (Note that 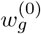 is by definition the number of leaves of genotype *g*, and is trivial to compute based on the input data.)
2. Iteratively construct the phylogeny by coalescing pairs of extant lineages, or by mutating one genotype to another according to the evolutionary path *g*_*i*_. Specifically: If a coalescence occurs at step *i*, choose a pair of extant lineages of genotype *g*_*i*_ uniformly at random to coalesce. If a mutation occurs, replace the single extant lineage of genotype *g*_*i*_ with a lineage of neighboring genotype *h*. As we show in Appendix A, there is only one such *h* that will lead to a valid tree. At this stage, do not specify the times of the coalescneces or mutations. After this step, we have constructed a RLMAT.
3. Sample each time interval Δ*t*_*i*_ as

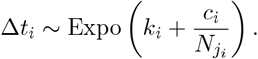

After doing so for all *i*, we have constructed a TLMAT.
4. Forget the mutation events to arrive at a timed, labeled tree.

See Table 1, rightmost two columns for an example of converting an evolutionary path back to a timed, labeled tree. By way of this procedure, together with the posterior distribution over evolutionary paths, we have established a Bayesian framework for inferring the posterior distribution of trees under the infinite sites mutation model that only requires updating a single sequence of discrete entries.

**Table 1:** Procedure for checking the validity of an evolutionary path and/or generating such a sequence. This procedure iterates over indices of an evolutionary path and updates the node weights following each coalescence and mutation. A coalescence may only occur between two extant lineages of the same genotype. A mutation may only occur in a genotype with one extant lineage, for which all but one adjacent subtree on the augmented perfect phylogeny has no extant lineages (shown grayed out). An example of a TLMAT corresponding to each evolutionary path is also shown. The unique genotypes on the augmented perfect phylogeny are labeled a, b, c, d, e.

| $i$ | Aug. Perf. Phylo. | $w^{(i)}$ | Evolutionary Path ($u$) | Tree |
| --- | --- | --- | --- | --- |
| 0 | | $(1, 0, 1, 0, 3)$ | | |
| 1 | | $(0, 1, 1, 0, 3)$ | | |
| 2 | | $(0, 1, 1, 0, 2)$ | | |
| 3 | | $(0, 1, 1, 0, 1)$ | | |
| 4 | | $(0, 1, 1, 1, 0)$ | | |
| 5 | | $(0, 0, 2, 1, 0)$ | | |
| 6 | | $(0, 0, 3, 0, 0)$ | | |
| 7 | | $(0, 0, 2, 0, 0)$ | | |
| 8 | | $(0, 0, 1, 0, 0)$ | | |

## 3 Inferring Trees and Evolutionary Parameters

### Overview

Given our theoretical framework for sampling trees under the infinite sites model as sequences of discrete entries, we now provide a Bayesian algorithm to infer not only the tree but also the effective population size trajectory. Recall that our goal is to sample from the posterior distribution

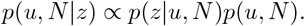

Breaking into cases based on whether *u* is compatible with *z*, and expanding the prior, the above reduces to

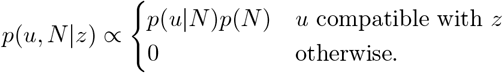

The term *p*(*u*|*N*) is derived in Section 2.6. For the prior on *N*, we assume the “Skyline” model (Section 3.2). We then propose a Gibbs scheme for sampling the posterior, i.e. we sample from

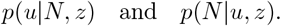

As we will see in Sections 3.1–3.2, our sampler for *p*(*u*|*N, z*) proposes relatively minor changes to the evolutionary path per iteration, whereas our sampler for *p*(*N*|*u, z*) proposes relatively major changes to the effective population size per iteration. Hence, in practice, we perform *M*_tree_ updates to *u* for every one update to *N*, and perform *M*_gibbs_ total updates to *N*, for some large *M*_tree_ and *M*_gibbs_. We call a series of *M*_tree_ updates to *u* followed by one update to *N* a *Gibbs cycle*. We then recover a timed phylogenetic tree from an evolutionary path *u* as discussed in Section 2.6, requiring only a single random sample.

### 3.1 Metropolis-Hastings Tree Inference

To explore the space of evolutionary paths *u* conditional on the effective population size trajectory *N*, we employ a simple Metropolis-Hastings scheme with a proposal distribution consisting of a single adjacent transposition move. Let *u* = (*g*_1_, …, *g*_*m*+*n−*1_). The adjacent transposition move proposes a new vector *u*^*′*^ by first selecting an index *i* ∈{1, …, *m* + *n –* 2} uniformly at random, and then swapping elements *g*_*i*_ and *g*_*i*+1_ in vector *u* to arrive at *u*^*′*^. The state is then updated according to the Metropolis-Hastings acceptance probability. We note that all evolutionary paths with nonzero posterior probability are in fact permutations of one another, though not all permutations are possible (shown in Appendices A and B). See Figure 2 for an example of an adjacent transposition move.

**Figure 2:**
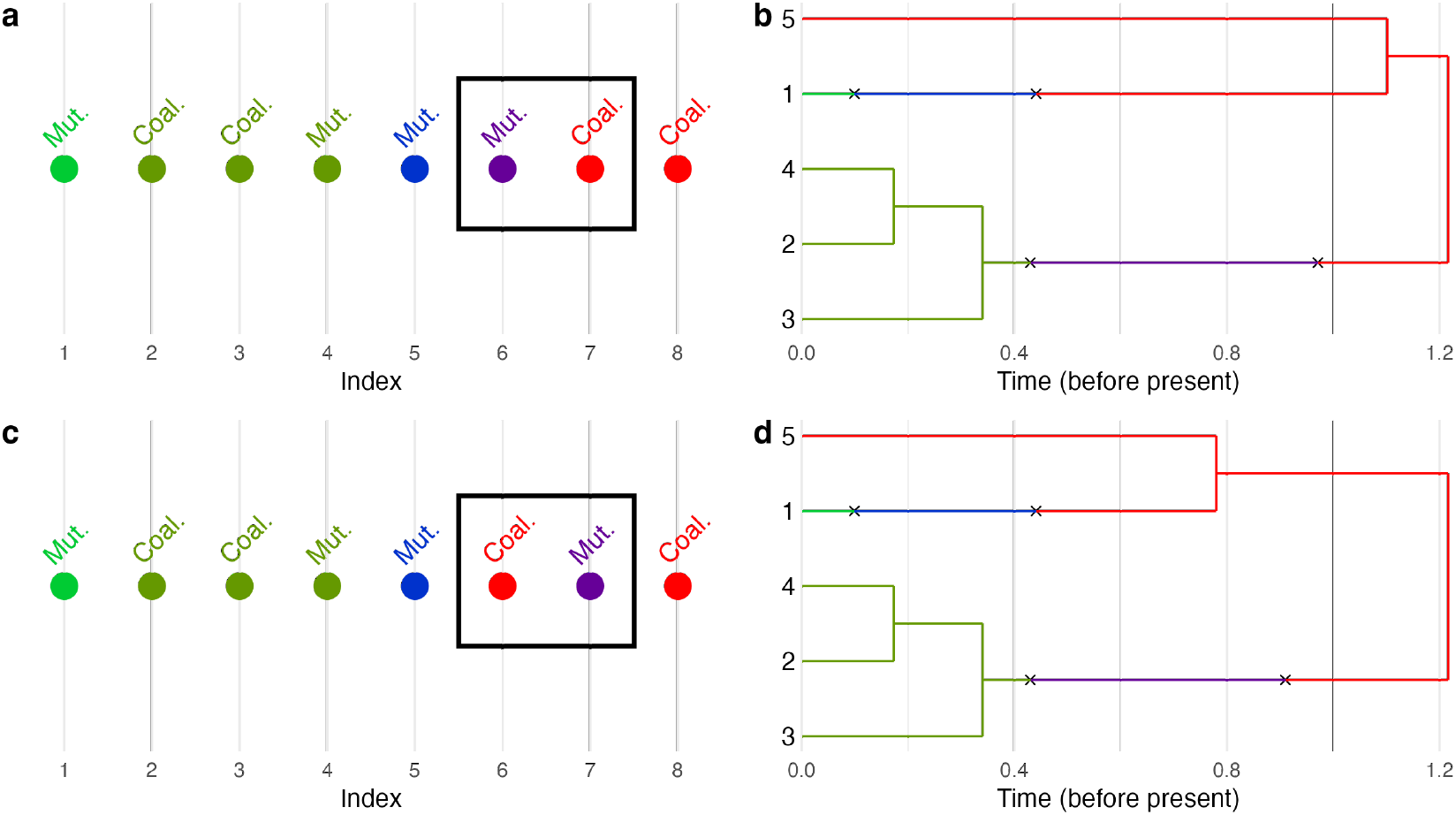
An adjacent transposition move from an evolutionary path (a) to (c), and examples of corresponding TLMATs (b) and (d), respectively. Recall that an evolutionary path does not store the times explicitly; other mutation and coalescent times are possible for (b) and (d).

Despite the simplicity of this move, substantial work is required to determine which proposals *u*^*′*^ are indeed compatible with *z* (Appendix A). We first establish a necessary condition for an evolutionary path to be compatible with a perfect phylogeny. For this, we define an *adjacent subtree of node g containing h* on the perfect phylogeny to consist of all nodes for which the path distance to *h* is less than the path distance to *g*. We then prove that a mutation in genotype *g* can occur only when (1) there is exactly one extant lineage of genotype *g*, (2) all but one adjacent subtree to *g* contains no genotypes with extant lineages, and (3) *g* is not the root genotype, if the root is specified by the user. An overview of the procedure for checking the validity of an evolutionary path is shown in Table 1. These conditions give us an *O*(1) runtime method of checking whether a proposed evolutionary path resulting from a transposition is valid and, if so, computing its posterior probability. Hence, the move may be repeated an enormous number of times while incurring minimal computational cost. Finally, in Appendix B, we prove that our Markov chain is irreducible.

### 3.2 Effective Population Size Inference

The previous section establishes a method for sampling evolutionary paths *u*, and hence trees, conditional on a fixed effective population size *N* and a known perfect phylogeny *z*. We now develop an inferential method for *N* conditional on *u* and *z*. The “Skyline” prior on *N* (Drummond et al., 2005) is given by

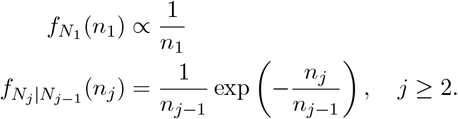

where 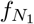 denotes the PDF of *N*_1_ and 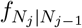 denotes the conditional PDF of *N*_*j*_ given *N*_*j−*1_. Note that the prior on *N*_1_, the Jeffreys prior, is improper. As we will see shortly, it is convenient to work instead with the log-transformed effective population sizes, which we denote by the vector *q*:

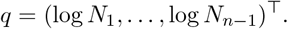

Applying this change of variables, the prior becomes

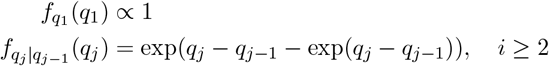

Once again, the prior on *q*_1_—equivalent to Unif(*−*∞, ∞)—is improper.

Now that we have defined the effective population size prior, we develop an efficient sampling strategy for *N* . This strategy is based on the “block update move” of the Skygrid model (Gill et al., 2013), but redesigned for Skyline without explicit representation of the coalescent times. Recall that, up to a constant of proportionality, the conditional density *p*(*N*|*z, u*) is given by:

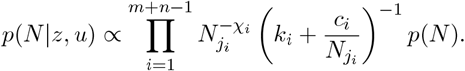

Writing out the prior *p*(*N*) and changing variables as *q*_*j*_ = log *N*_*j*_, we obtain:

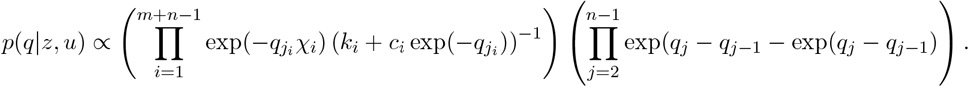

To simplify this expression, first, note that *j*_*i*_ only takes values 1, 2, …, *n −* 1. For each *j* ∈ *{*1, …, *n −* 1*}*, there is exactly one value *i* such that *j*_*i*_ = *j* and *χ*_*i*_ = 1; for all other *i* with *j*_*i*_ = *j*, we have *χ*_*i*_ = 0. Moreover, if *j*_*i*_ = *j*_*i*_*′* then *k*_*i*_ = *k*_*i*_*′* and *c*_*i*_ = *c*_*i*_*′* . Hence, let *m*_*j*_ := |{*i* : *j*_*i*_ = *j*}|, let *k*_*j*_ := *k*_*i*_ for *i* such that *j*_*i*_ = *j*, and analogously for *c*_*j*_. Therefore,

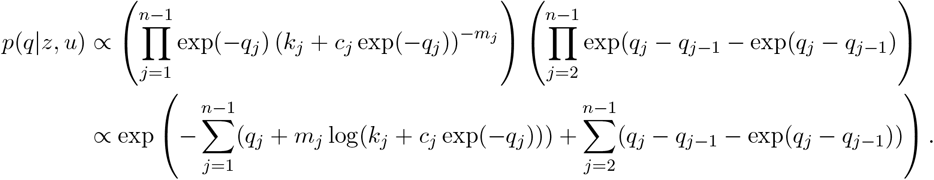

To sample from this conditional density, we generate a proposal using Laplace approximation (see, e.g., MacKay (2003)). We use numerical methods to obtain the mode of *p*(*q*|*z, u*), which we set equal to the mean of our approximating Gaussian density (as the mean and mode are equal in a Multivariate Normal distribution). We then use a Taylor expansion at this mode to compute the variance-covariance matrix for this density, which allows us to approximate *p*(*q*|*z, u*) to second-order accuracy. Finally, we propose *q* from this Gaussian, and accept or reject based on the MetropolisHastings condition. The details are left to Appendix C.

## 4 Results

### 4.1 Inferring Trees Alone

First, we validated and benchmarked our approach by reconstructing the evolutionary histories of synthetic datasets. Synthetic datasets were generated by sampling trees from the coalescent model, overlaying mutations on each branch according to a Poisson process, and generating sequences in accordance with the infinite sites assumption. We simulated trees with 32, 128, and 512 taxa, and with constant effective population sizes (relative to mutation rate *µ* = 1) of 4, 16, and 64, making for 9 total datasets. To assess the accuracy and statistical efficiency of the tree topology inference scheme alone, as opposed to the joint inference of the tree and effective population size, we instructed `inPhynite` not to infer the effective population size, and left it fixed at the true value (unknown effective population size trajectories are addressed in Sections 4.2–4.3). We performed 100,000,000 Metropolis Hastings tree moves for each synthetic dataset.

As a baseline, we compared our results to `BEAST 2`. While `BEAST 2` does not offer the infinite sites mutation model, this model can be well approximated by the Jukes-Cantor model (Jukes and Cantor, 1969) when the number of polymorphic sites across the input dataset is small compared to the genome length. To achieve this, we simulated sequence datasets from our perfect phylogenies with genome length 100,000. We ran `BEAST 2` with a total of 1,000,000 iterations, again fixing the effective population size (and mutation rate) at its true value. For both methods, we then extracted 1,000 posterior trees and discarded the first 20% as burn-in.

We observed strong concordance between `inPhynite` and `BEAST 2` in terms of the distribution of tree topologies and coalescent times, and dramatically greater statistical efficiency for `inPhynite` as compared to `BEAST 2`. We plotted the distributions of the inferred time of most recent common ancestor (TMRCA), inferred Robinson-Foulds distance to the true tree (Robinson and Foulds, 1981), Frobeinus-norm distance based on *F* matrices (a lossless representation of unlabeled trees proposed by Kim et al. (2020)), and entry-wise bias and variance of the *F* matrix (see Figure 3 and Appendix E, Figures S1–S2 for detailed comparisons on the datasets with 32 taxa). Posterior distributions were nearly identical, with minor discrepancies attributable to sample size. To assess statistical efficiency, we computed the overall effective sample size per unit runtime using an 2.6 GHz 6-Core Intel Core i7 processor, where overall effective sample size was taken to be the minimum of the effective sample sizes of the coalescent times and the *F* matrix entries (see Table 2). `inPhynite` consistently outperformed `BEAST 2` by this metric, with particularly drastic speedup on larger datasets—achieving over 225 times greater statistical efficiency on the dataset with 512 taxa and effective population size 64.

**Table 2:** Performance comparison between `inPhynite` and `BEAST 2` for fixed effective population size. Numbers in bold indicate superior efficiency.

| Taxa | $N$ | Runtime (sec) | | Overall ESS | | Efficiency (ESS/sec) | | Speedup |
| --- | --- | --- | --- | --- | --- | --- | --- | --- |
|  |  | inPhynite | BEAST 2 | inPhynite | BEAST 2 | inPhynite | BEAST 2 |  |
| 32 | 4 | 3.70 | 10.5 | 207 | 437 | **55.8** | 41.7 | **1.34** |
| 32 | 16 | 2.89 | 11.8 | 546 | 431 | **189** | 36.4 | **5.19** |
| 32 | 64 | 2.72 | 12.5 | 390 | 453 | **144** | 36.2 | **3.96** |
| 128 | 4 | 2.95 | 32.0 | 392 | 93.4 | **133** | 2.92 | **45.5** |
| 128 | 16 | 3.69 | 35.6 | 324 | 138 | **87.8** | 3.87 | **22.7** |
| 128 | 64 | 3.52 | 40.3 | 285 | 95.6 | **81.0** | 2.37 | **34.2** |
| 512 | 4 | 4.04 | 184 | 207 | 57.0 | **51.1** | 0.310 | **165** |
| 512 | 16 | 5.09 | 167 | 189 | 34.6 | **37.2** | 0.207 | **180** |
| 512 | 64 | 5.63 | 209 | 161 | 26.4 | **28.5** | 0.126 | **226** |

**Figure 3:**
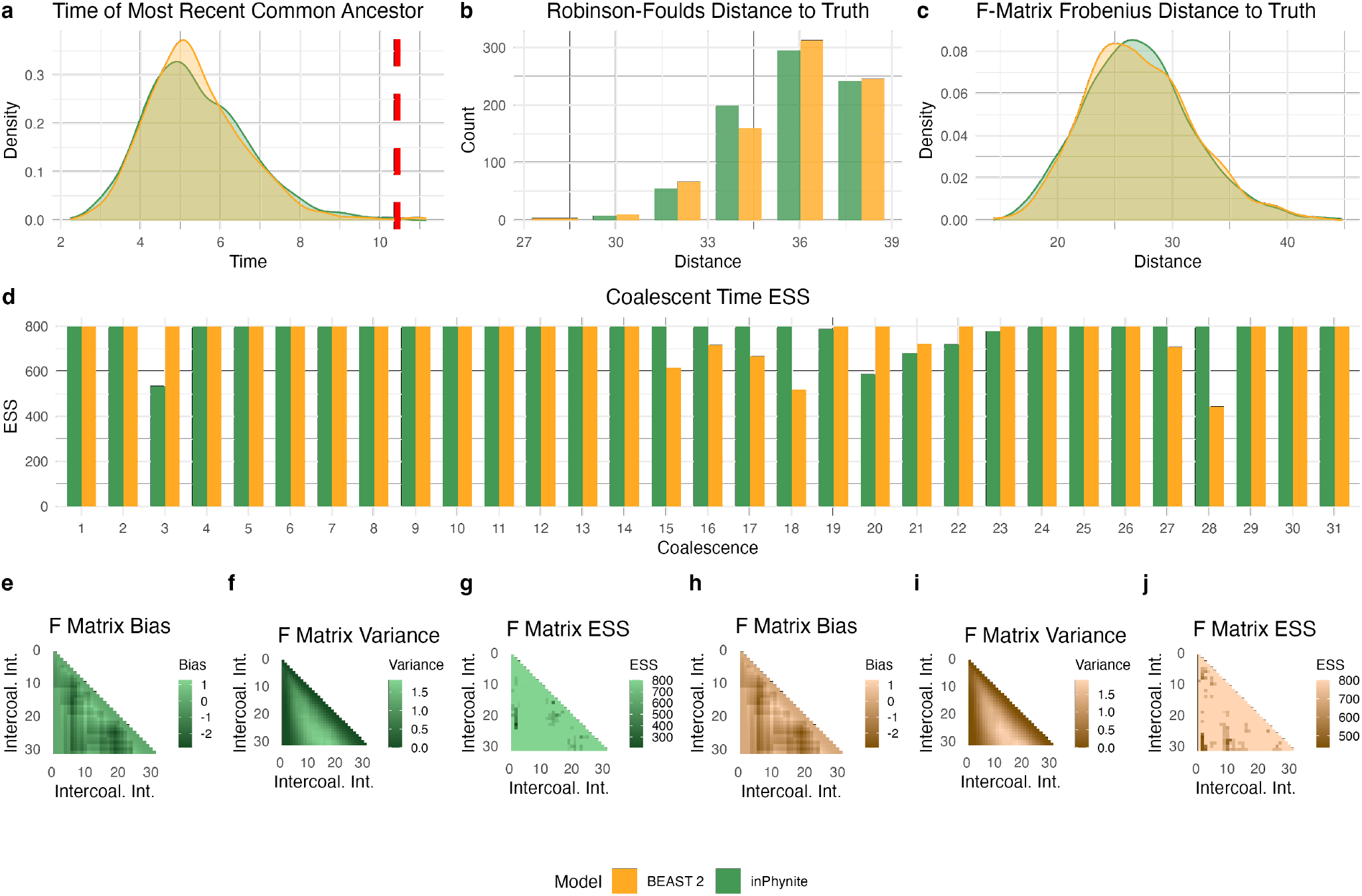
Performance comparison of `inPhynite` (green) to `BEAST 2` (orange) for 32 taxa and effective population size *N*_*j*_ = 4 for each intercoalescent interval *j*. (a) Posterior distribution of the time of most recent common ancestor; (b) Robinson-Foulds distance to the true tree; (c) Frobenius norm of the difference between true and inferred *F* matrices; (d) Effective sample size for each coalescent time; (e)–(j) bias, variance, and entry-wise ESS of the *F* matrices of the posterior samples.

### 4.2 Inferring Trees and Effective Population Sizes

We next compared `inPhynite` to `BEAST 2` in the scenario where the effective population size varied over time, using synthetic datasets with 40 taxa each. We considered three types of effective population size trajectories: constant, exponential growth, and exponential decay, and approximated each as a piecewise-constant function as to match the Skyline model. Specifically, for *t* measured in time units since the present, the effective population size at the beginning of each intercoalescent interval is given by the function

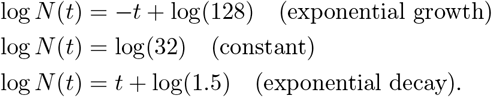

For `inPhynite`, we performed 100,000 total Gibbs cycles on each dataset (*M*_gibbs_ = 10^5^). We updated the tree 10,000 times per update of the effective population size trajectory (*M*_tree_ = 10^4^). We ran `BEAST 2` under the Coalescent Bayesian Skyline model (Drummond et al., 2005) with 100,000,000 iterations. As before, we sampled 1,000 total trees and effective population size trajectories, and discarded the first 20% of the samples as burn-in.

Once again, we observed strong concordance between the inferences of the two methods, and an improvement in statistical efficiency for `inPhynite` over `BEAST 2`. In addition to the posterior distributions we discussed for fixed *N*, we also visualized the posterior distribution of the effective population size trajectory, which was nearly identical between the two methods (see Figure 4 and Appendix F, Figures S3–S4). We calculated overall effective sample size for both methods to be the minimum over effective sample sizes based on the coalescent times, *F* matrix entries, and effective population sizes on each intercoalescent interval (see Table 3). We observed a significantly greater speedup here as compared to on similarly sized datasets in the previous analysis (i.e. with fixed effective population size), indicating that `inPhynite`‘s method for sampling *N* is more efficient than that of `BEAST 2`.

**Table 3:** Performance comparison between `inPhynite` and `BEAST 2` for inferred effective population size. Numbers in bold indicate superior efficiency.

| Scenario | Runtime (sec) |  | Overall ESS |  | Efficiency (ESS/sec) |  | Speedup |
| --- | --- | --- | --- | --- | --- | --- | --- |
|  | inPhynite | BEAST 2 | inPhynite | BEAST 2 | inPhynite | BEAST 2 |  |
| Growth | 45.8 | 1511 | 551.4 | 235.4 | **12.0** | 0.156 | **77.2** |
| Constant | 38.7 | 1403 | 372.8 | 240.6 | **9.63** | 0.171 | **56.2** |
| Decay | 40.6 | 1292 | 316.1 | 173.1 | **7.79** | 0.134 | **58.1** |

**Figure 4:**
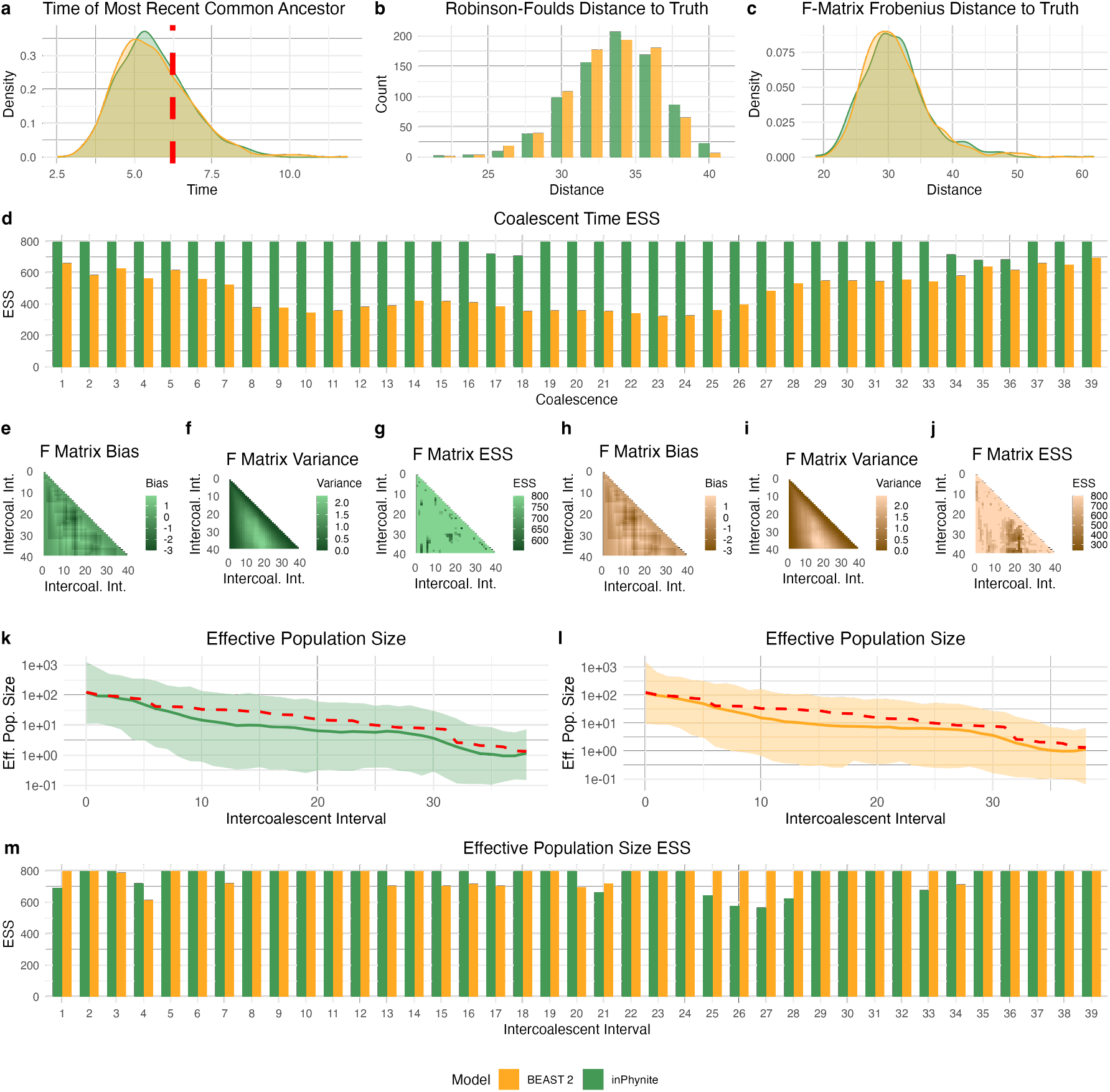
`inPhynite` and `BEAST 2` inferences and performance for an exponentially growing effective population size trajectory. (a) Posterior distribution of the time of most recent common ancestor; (b) Robinson-Foulds distance to the true tree; (c) Frobenius norm of the difference between true and inferred *F* matrices; (d) Effective sample size for each coalescent time; (e)–(j) bias, variance, and entry-wise ESS of the *F* matrices of the posterior samples; (k)–(l) Posterior distribution of the effective population size trajectory; (m) Effective sample size of the effective population size on each intercoalescent interval.

### 4.3 Real Data Application

We used `inPhynite` to learn about the genealogy and effective population size of various human populations over time, as well as to compare our method to `BEAST 2` using real data. We analyzed mitochondrial data samples collected from four human groups as part of the 1,000 Genomes Project: Utah residents with Northern/Western European ancestry (CEU), Finnish people in Finland (FIN), Peruvians in Lima (PEL), and Yoruba in Ibadan, Nigeria (YRI) (1000 Genomes Project Consortium, 2015). We retained only the coding region of each sequence, spanning sites 576–16,024 (Anderson et al., 1981; Andrews et al., 1999). We ran `BEAST 2` on the resulting alignments using a mutation rate 1.3 × 10^*−*8^ substitutions per site per year (Palacios et al., 2019). For `inPhynite`, we masked a further 25 of 336, 15 of 232, 25 of 311, and 54 of 489 polymorphic sites, respectively, that were not compatible with the infinite sites model. Due to this masking, we used a mutation rate of 1.3 × 10^*−*8^ multiplied by the proportion of polymorphic sites compatible with the infinite sites model. Again, for both methods, we extracted 1,000 posterior trees and effective population size trajectories, and discarded the first 20% as burn-in.

Inferred effective population size trajectories generally agreed between the two methods, though we observed some discrepancies that can be attributed to the polymorphic sites masked by `inPhynite` and not by `BEAST 2` (see Figure 5). Nonetheless, these trajectories showed the same overall trends, and 95% posterior credible intervals consistently overlapped between the methods. In accordance with our synthetic data studies, `inPhynite` once again achieved higher statistical efficiency than `BEAST 2`. These results suggest that for human mitochondrial DNA datasets, only a modest amount of information about the effective population size is lost by masking sites incompatible with the infinite sites model.

**Figure 5:**
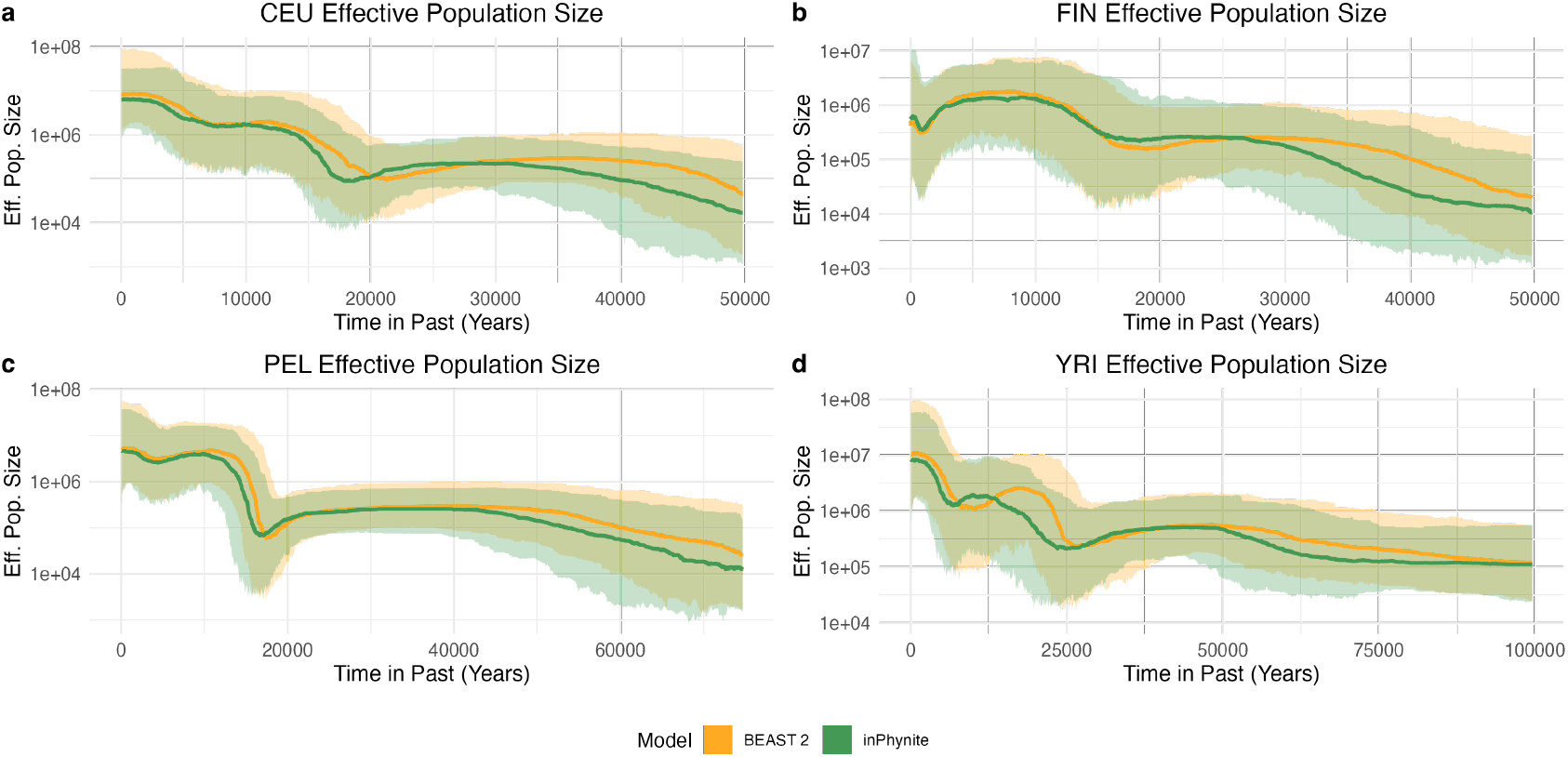
Effective population size of (a) Utah residents with Northern/Western European ancestry (CEU), (b) Finnish people in Finland (FIN), (c) Peruvians in Lima (PEL), and (d) Yoruba in Ibadan, Nigeria (YRI), as inferred by `inPhynite` and `BEAST 2`.

## 5 Discussion

Bayesian phylogenetics algorithms face computational challenges when dealing with large datasets and variable effective population size trajectories. These challenges arise largely due to the enormous cardinality of timed, labeled trees, causing MCMC methods to mix slowly. Under the infinite sites model, however, we show that trees may instead be sampled at the far lower resolution of *evolutionary paths*, i.e. sequences of the genotypes in which each coalescence or mutation occurs. By developing simple yet efficient MCMC moves to explore the space of evolutionary paths and effective population size trajectories, we achieve rapid convergence on both real and synthetic datasets with our method, `inPhynite`.

Considering the diversity of models employed by Bayesian phylogeneticists, we envision numerous future developments to `inPhynite` that would make it compatible with a wider range of datasets. In terms of effective population size models, we plan to incorporate the Skygrid prior (Gill et al., 2013), in which the effective population size is constant on prespecified time intervals as opposed to intercoalescent intervals. To do so will require storing the times of each coalescence and mutation explicitly. While this modification unfortunately doubles the size of the evolutionary parameter space, and will likely lead to slower convergence and higher cost per iteration, our main idea from Section 2.6—the representation of trees as evolutionary paths—remains possible, so we would still expect gains in statistical efficiency as compared to existing methods. We also aim to incorporate heterochronous sampling into future versions, which will be useful for analyzing datasets with significant heterogenetity in sample collection date relative to total evolutionary time (e.g. ancient DNA, infectious disease outbreaks).

We further plan to explore inference algorithms beyond MCMC for Bayesian infinite sites phylogenetic inference. Considering the fact that the posterior *p*(*u*|*z, N*) can be expressed as the distribution of a Markov chain with state space given by the number of extant lineages of each genotype following each coalescence or mutation event, Sequential Monte Carlo is a natural choice of inference algorithm (Chopin and Papaspiliopoulos, 2020). This approach offers the major practical advantage that tree proposals are highly parallelizable, enabling runtime improvements via multithreading. We provide the details of an SMC approach in Appendix D. Even with parallel processing, however, we were unable to match the statistical efficiency of the Metropolis-Hastings scheme for larger datasets (≈500 total mutations and coalescences) due to severe particle degeneracy. Nonetheless, we hope to revisit this idea in subsequent work.

Although `inPhynite` is designed for handling genomic datasets compatible with the infinite sites mutation model, it could reasonably be applied to any set of taxa in which one perfect phylogeny is overwhelmingly more probable than all others. In Bayesian phylogenetic analysis of viral outbreaks, for instance, it is common for many posterior samples to share the same underlying perfect phylogeny, and for that phylogeny to be parsimonious or approximately parsimonious (Varilly et al., 2025). In such scenarios, a parsimonious perfect phylogeny could be input to `inPhynite` to sample an approximate posterior distribution rapidly. Although finding a parsimony tree from a dataset of sequences is NP-hard, we are exploring possible algorithms that can do so in polynomial time if repeated mutations at a site on the genome are sufficiently rare. Such algorithms can be used to check whether a dataset could reasonably be modeled as having a fixed underlying perfect phylogeny, and, if so, provide said phylogeny as input to `inPhynite`.

Generalizing `inPhynite` to the finite-sites context, i.e. relaxing the assumption of a fixed perfect phylogeny, poses a far greater challenge. To do so, we must modify the definition of an evolutionary path to record the resulting new genotype on the tree following each mutation event. The difficulty here is that unlike in the infinite sites case, *any* genotype is theoretically possible, though some are much more likely to appear than others. Moreover, the total number of mutations on the tree is no longer a fixed quantity, meaning that evolutionary paths must now have variable length.

Though there are many possible avenues for expansion, `inPhynite` offers a promising novel framework for phylogenetic tree reconstruction. We aim for our method of representing the latent tree space as a sequence of genotypes in which each coalescence or mutation occurs to serve as the basis for efficient inference algorithms under a wide variety of different models. With `inPhynite`, we hope to push the size and scope of dataset for which Bayesian phylogenetic inference is possible.

## Acknowledgements

The authors acknowledge I.H. Goldstein, G.R. Mazzeo, P.C. Sabeti, and P. Varilly for their valuable feedback on the manuscript.

## Code and Data Availability

`inPhynite` is available to the public as a command-line tool at github.com/ispecht/inPhynite. It is written in `C++`. Data used in this study are available to the public through the 1,000 Genomes Project (1000 Genomes Project Consortium, 2015).

## Supplementary Material

### A Proof of Conditions for a Valid Evolutionary Path

Here, we present and justify the conditions necessary for an evolutionary path (*u*) to be compatible with a perfect phylogeny, as well as develop an efficient procedure for checking whether an adjacent transposition Metropolis-Hastings move preserves the validity of an evolutionary path. We first develop a method for keeping track of the number of extant lineages of each genotype at the time of each coalescence or mutation. We propose a structure called a (*χ, p*) sequence (for reasons that will soon become clear) to record the coalescences and mutations on a phylogeny, which is slightly different than an evolutionary path in that a (*χ, p*) sequence explicitly stores both the identity (coalescence or mutation) of each event as well as the genotype(s) involved in each event—one genotype for a coalescence, and two genotypes for a mutation. From there, we define what it means for a (*χ, p*) sequence to be valid, i.e. compatible with a perfect phylogeny, and prove a key necessary condition for validity. Finally, we show that this notion of validity not only extends from (*χ, p*) sequences to evolutionary paths, but is also straightforward to compute.

Let *z* be an augmented perfect phylogeny. Let the nodes of *z* be denoted by a set *V* . Recall that the nodes of *z* correspond to the set of genotypes observed in a dataset, as well as all genotypes that must have occurred at some point in the evolutionary past. For that reason, we will use the words “nodes” and “genotypes” interchangeably moving forward. Let *E* denote the set of edges of *z*. Recall that the edges of *z* connect pairs of genotypes in which one evolved from the other by way of a single mutation. Finally, let *R* ∈ *V* denote the optional root node of *z*, i.e. the genotype ancestral to all of the samples. If *R* is not specified by the user, we use the notation *R* = ∅.

Next, we formalize the notions of coalescence and mutation, and establish a method for recording the number of extant lineages of each genotype before/after each coalescence and mutation event. To do so, first fix any enumeration of the set *V*, such that we may refer to the “*j*th genotype” unambiguously for 1 ≤ *j* ≤ |*V*| . Let *w*^(0)^ be a tuple of non-negative integers of length *V*, where the *j*th entry of *w*^(0)^ equals the number of sampled tips of genotype *j*. For the example in Figure 1d, the tuple *w*^(0)^ would be given by

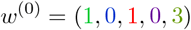

where the fixed enumeration of the genotypes in *V*, represented as colors in the augmented perfect phylogeny in Figure 1e, is

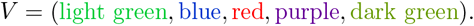

Now that we have a method for recording the number of extant lineages of each genotype, we may view coalescences and mutations as *actions* taken to alter the genotype counts in one of two ways. If a coalescence occurs in a certain genotype, the number of extant lineages of that genotype decreases by one—provided there were two or more to begin with. If a mutation occurs in a certain genotype, the number of extant lineages of that genotype decreases by one, and the number of extant lineages of a neighboring genotype on the perfect phylogeny increases by one. A mutation can only occur in a genotype with one or more extant lineage to begin with.

To formalize this idea, we introduce variables *p*_*i*_ ∈ *V* ⊔ (*V × V*) that record either a single genotype in which a coalescence occurs, or the ordered pair of genotypes in which a mutation occurs (the first of the pair being the genotype in which we decrease the number of extant lineages by one, and the second being the neighboring genotype in which we increase the number of extant lineages by one).

The index *i* here ranges from 1 to *m* + *n* − 1, so that we have one variable *p*_*i*_ corresponding to each coalescence and mutation event. Like in the main text, we define *χ*_*i*_ to be the indicator of whether the *i*th event is a coalescence, i.e. *χ*_*i*_ = 𝟙(*p*_*i*_ ∈ *V*). For the example given in Figure 1d, the *χ*_*i*_’s and *p*_*i*_’s are given by

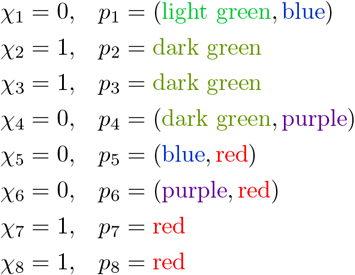

Now that we have defined whether the *i*th action is a coalescence or mutation, and have further specified which genotype(s) are affected by the action, we introduce a series of tuples *w*^(1)^, …, *w*^(*m*+*n−*1)^ to record the number of extant lineages of each genotype after the *i*th action, for *i* = 1, 2, …, *m* + *n ™* 1. These are defined recursively as follows: First, if *χ*_*i*_ = 1, i.e. the *i*th action is a coalescence, then

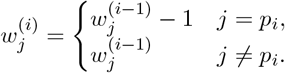

where 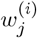 denotes the *j*th entry of *w*^(*i*)^. Now, if *χ*_*i*_ = 0, i.e. the *i*th action is a mutation, write *p*_*i*_ = (*g*_*i*_, *h*_*i*_) for some *g*_*i*_, *h*_*i*_ ∈ *V* . Then

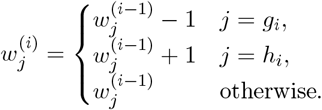

For the example given in Figure 1d, the indicators *χ*_*i*_, the sequence *w*^(0)^, *w*^(1)^, …, *w*^(*m*+*n−*1)^ is given by:

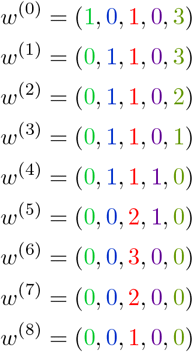

We refer to the coalescence indicators *χ*_1_, …, *χ*_*m*+*n−*1_, together with the genotype(s) *p*_1_, …, *p*_*m*+*n−*1_ in which each action occurs, as a (*χ, p*) *sequence*. As we just saw, the tuples *w*^(0)^, *w*^(1)^, …, *w*^(*m*+*n−*1)^ are deterministic functions of (*χ, p*). However, not all possible sequences (*χ, p*) correspond to valid phylogenetic trees: for example, some choices of (*χ, p*) may lead to tuples *w*^(*i*)^ with negative entries, and having a negative number of extant lineages of a genotype is nonsensical. For this reason, we introduce the concept of a *valid* (*χ, p*) *sequence*, defined as follows:

#### Definition 1.

Consider a (*χ, p*) sequence, and let *w*^(0)^, *w*^(1)^, …, *w*^(*m*+*n−*1)^ be defined based on (*χ, p*) as above. We say that (*χ, p*) is *valid* if the following three axioms hold:

1. If *χ*_*i*_ = 1, then 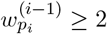.
2. If *χ*_*i*_ = 0, let *p*_*i*_ = (*g, h*). Then 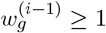 and *{g, h}* ∈ *E*.
3. If *R* ≠ ∅, then

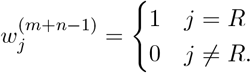

Intuitively, these axioms state that:

1. A coalescence can only occur between two extant lineages of the same genotype.
2. A mutation converts an extant lineage of one genotype into a lineage of an adjacent genotype on an augmented perfect phyloegny.
3. After all coalescences and mutations have occurred, we are left with one extant lineage of the root genotype.

With this framework in place, we are nearly ready to state and to prove a necessary condition for validity of a (*χ, p*) sequence. Before doing so, we introduce a few useful definitions relating to trees:

#### Definition 2

Let *z* = (*V, E*) be a tree and let *g, h* ∈ *V* . The *path distance d*(*g, h*) from *g* to *h* is defined as the minimum integer *k* such that there exist *g*_0_, *g*_1_, … *g*_*k*_ with *g*_0_ = *g, g*_*k*_ = *h*, and *{g*_*i−*1_, *g*_*i*_*}* ∈ *E* for all 1 ≤ *i* ≤ *k*. If *g* = *h*, we set *d*(*g, h*) = 0.

Using this distance *d*, we define an *adjacent subtree* as follows:

#### Definition 3

Let *z* = (*V, E*) be a tree and let *g, h* ∈ *V* . The *subtree containing h adjacent to g*, denoted *z*_*h*:*g*_, is the subtree defined by

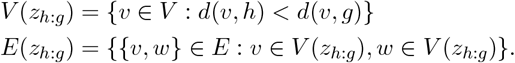

It is also helpful to define the notion of a subtree containing no extant lineages:

#### Definition 4

Let *z, V, E, g*, and *h* be as above, and let *w* be a tuple of non-negative integers of length |*V*| . Let *w*_*j*_ denote the *j*th entry of *w*. We say that *w vanishes on z*_*h*:*g*_ if for all *j* ∈ *V* (*z*_*h*:*g*_), we have *w*_*j*_ = 0.

We may now state our main result, which will imply a tractable method for checking the validity of an evolutionary path:

#### Proposition 1

Let *z* be an augmented perfect phylogeny and suppose (*χ, p*) is valid. Construct *w*^(0)^, *w*^(1)^, …, *w*^(*m*+*n−*1)^ based on (*χ, p*) as before. Consider *i* with *χ*_*i*_ = 0, i.e. action *i* is a mutation, and write *p*_*i*_ = (*g, h*). Then for all such *i*,

1.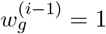.

2. For all *h*^*′*^ ≠ *h*, if *{g, h*^*′*^*}* ∈ *E* then *w*^(*i−*1)^ vanishes on 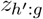.

3. *g* ≠ *R*, if *R* ≠ ∅.

Intuitively, these conditions mean:

1. A mutation can only happen in a genotype for which there is one extant lineage.
2. A mutation from genotype *g* to *h* can only happen if for every other genotype *h*^*′*^ adjacent to *g* on the perfect phylogeny, the subtree adjacent to *g* containing *h*^*′*^ has no extant lineages.
3. If the root is specified, a mutation cannot happen in the root genotype.

The proof is facilitated by the following lemma:

#### Lemma 1

Let (*χ, p*) be valid, and construct *w*^(0)^, *w*^(1)^, …, *w*^(*m*+*n−*1)^ based on (*χ, p*) as before. Fix some *k* ≥ 0, and select any edge *{g, h}* ∈ *E*. Suppose that *w*^(*k*)^ does not vanish on *z*_*h*:*g*_ and *w*^(*k*)^ also does not vanish on *z*_*g*:*h*_. Then for some *k*^*′*^ *> k*, we have either *p*_*k*_*′* = (*g, h*) or *p*_*k*_*′* = (*h, g*).

*Proof of Lemma*. Suppose for contradiction that for all *k*^*′*^ *> k, p*_*k*_*′* (*g, h*) and *p*_*k*_*′* (*h, g*). We claim that each *w*^(*k*^*′*^)^ vanishes neither on *z*_*h*:*g*_ nor *z*_*g*:*h*_. We proceed by induction. The base case *k*^*′*^ = *k* is given in the statement of the lemma. For the inductive step, suppose that *w*^(*j*)^ vanishes neither on *z*_*h*:*g*_ nor *z*_*g*:*h*_. First suppose the (*j* + 1)st action is a coalescence, i.e. *χ*_*j*+1_ = 1. By

Definition 3, axiom 1, we have 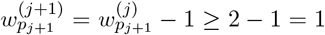, so *w*^(*j*+1)^ does not vanish at *p* . Moreover *w*^(*j*+1)^ agrees with *w*^(*j*)^ everywhere except for *p*_*j*+1_; thus, *w*^(*j*+1)^ does not vanish on *z*_*h*:*g*_ or *z*_*g*:*h*_.

Now suppose *χ*_*j*+1_ = 0, i.e. the (*j* + 1)st action is a mutation, and write *p*_*j*+1_ = (*g*_*j*+1_, *h*_*j*+1_). By assumption that (*g*_*j*+1_, *h*_*j*+1_) ∈*/* (*g, h*), (*h, g*), we have either *g*_*j*+1_, *h*_*j*+1_ ∈ *V* (*z*_*h*:*g*_) or *g*_*j*+1_, *h*_*j*+1_ ∈ *V* (*z*_*g*:*h*_). Without loss of generality, assume *g*_*j*+1_, *h*_*j*+1_ ∈ *V* (*z*_*h*:*g*_). Since the (*j* + 1)st action is a mutation that only affects the counts of the numbers of extant lineages on *V* (*z*_*h*:*g*_), and since mutations do not change the *total* number of extant lineages, we obtain

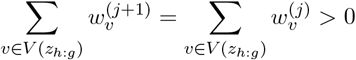

by the inductive hypothesis. This completes the proof by induction. We conclude that *w*^(*m*+*n−*1)^ in particular vanishes neither on *z*_*h*:*g*_ nor *z*_*g*:*h*_, contradicting Definition 3, axiom 3 and thus proving the lemma.

This lemma, combined with a counting argument, allows us to prove Proposition 1.

*Proof of Proposition*. We begin with some key observations. In order for (*χ, p*) to be valid, exactly *n −* 1 of the actions must be coalescences. To see this, note that if *χ*_*i*_ = 1, then

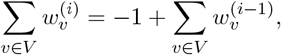

and if the *i*th action is a mutation, then

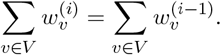

Since

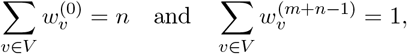

it follows that *χ*_*i*_ = 1 exactly *n* − 1 times. Now, consider the set of all *i* such that *χ*_*i*_ = 0, i.e the *i*th action is a mutation. We claim that for every edge {*g, h*} ∈ *E*, we can find some *i* such that *p*_*i*_ = (*g, h*) or *p*_*i*_ = (*h, g*). This follows immediately from Lemma 1 applied with *k* = 0, noting that *w*^(0)^ is nonvanishing on *V* (*z*_*h*:*g*_) and *V* (*z*_*g*:*h*_) because *w*^(0)^ is nonvanishing on all leaves of *z*. Therefore, the set {*p*_*i*_ : *i* is a mutation} is in bijection with *E*: every edge in *E* appears as the set of elements of *p*_*i*_ for exactly one *i*.

This bijection allows us to prove all parts of Proposition 1. Recall that for event *i* with *χ*_*i*_ = 0, we write *p* = (*g, h*) for some *g, h* ∈ *V* . For part (1), suppose for contradiction that 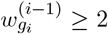. Then 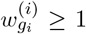 and 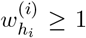. Since *w*^(*i*)^ is nonvanishing on 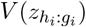 and 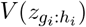, by the lemma there exists *j > i* such that *p*_*j*_ = (*g*_*i*_, *h*_*i*_) or *p*_*j*_ = (*h*_*i*_, *g*_*i*_). This contradicts the bijection between mutations and edges *E* of *z*.

For (2), suppose for contradiction there exists *h*^*′*^ *= h*_*i*_ such that *{g*_*i*_, *h*^*′*^*}* ∈ *E* and *w*^(*i−*1)^ is nonvanishing on 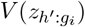. Since *w*^(*i*)^ agrees with *w*^(*i−*1)^ except at *g*_*i*_ and *h*_*i*_, and since neither *g*_*i*_ nor *h*_*i*_ is an element of 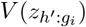, we know that *w*^(*i*)^ is nonvanishing on 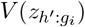. Moreover, since 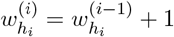, we know that *w*^(*i*)^ is nonvanishing on 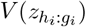. Since 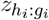 is a subtree of 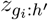, it follows that *w*^(*i*)^ is nonvanishing on 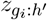. Thus, applying the lemma with *g* = *g*_*i*_ and *h* = *h*^*′*^ implies there exists *j > i* such that *p*_*j*_ = (*g*_*i*_, *h*^*′*^) or *p*_*j*_ = (*h*^*′*^, *g*_*i*_). As before, this contradicts the bijection between mutations and edges.

For (3), suppose for contradiction that *g*_*i*_ = *R* with *R* = ∅. Assume (1) and (2) hold, else we arrive at a contradiction immediately. It follows that *w*^(*i*)^ vanishes on 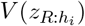. We claim that for all *j > i, w*^(*j*)^ must also vanish on 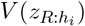. To see this, note that if *χ*_*j*_ = 1, it is impossible to have 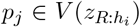 by Definition 1, axiom 1. If *χ*_*j*_ = 0 with *p*_*j*_ = (*g*_*j*_, *h*_*j*_), it is impossible to have 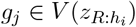, else we would contradict Definition 1, axiom 2. Moreover, we also cannot have 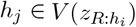: the only possible pair (*g*_*j*_, *h*_*j*_) with 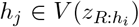 and 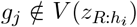 is (*g*_*j*_, *h*_*j*_) = (*h*_*i*_, *R*), and this mutation cannot occur again by the bijection between mutations and edges. We conclude that *w*^(*m*+*n−*1)^ vanishes on 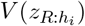, implying 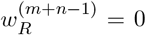 and thus contradicting Definition 1, axiom 3. The proof is complete.

What we have just proven establishes a necessary condition for a sequence of coalescences and mutations to be valid. Observe that, per terminology established in Section 2.6, a sequence of coalescences and mutations (*χ, p*) implies a unique evolutionary path:

#### Definition 5

The *implied evolutionary path* corresponding to (*χ, p*) is defined by

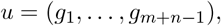

with

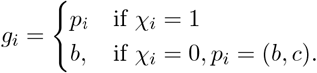

This idea allows us to extend the notion of validity to evolutionary paths:

#### Definition 6

Let *u* be an evolutionary path. Suppose there exists a valid (*χ, p*) sequence whose implied evolutionary path equals *u*. Then *u* is a *valid evolutionary path*.

We claim that if such an (*χ, p*) exists, then it is unique:

**Corollary 1**. Let *u* = (*g*_1_, …, *g*_*m*+*n−*1_) be an evolutionary path, let *z* and *w*^(0)^ be fixed and known, and suppose there exists a *valid* sequence of coalescences and mutations (*χ, p*) such that the implied evolutionary path of (*χ, p*) equals *u*. Then (*χ, p*) is unique.

*Proof*. Define *w*^(0)^ as before. For a given *u, w*^(0)^, and *z*, we construct (*χ, p*) and *w*^(1)^, …, *w*^(*m*+*n−*1)^, iteratively, showing that for each *i*, there is only one possible choice of *χ*_*i*_ and *p*_*i*_, and hence only one possible choice of *w*^(*i*)^. First, suppose 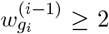. We claim *χ*_*i*_ = 1, i.e. the *i*th action in the sequence of coalescences and mutations must be a coalescence. To see this, suppose for contradiction that *χ*_*i*_ = 0, i.e. the *i*th action is a mutation. Write *p*_*i*_ = (*a, b*). Since *w*^(*i*)^ is nonvanishing on *z*_*a*:*b*_ and *z*_*b*:*a*_, Lemma 1 implies that *p*_*j*_ = (*a, b*) or *p*_*j*_ = (*b, a*) for some *j > i*, contradicting the bijection between edges of *z* and mutations established in the proof of Proposition 1. Hence, 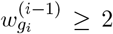 implies the *i*th action is a coalescence, so we set *χ*_*i*_ = 1, *p*_*i*_ = *g*_*i*_, and

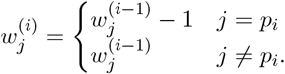

Now, suppose 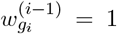 (the only remaining case, given the definition of a valid sequence of coalescences and mutations). It follows that *χ*_*i*_ = 0, so write *p*_*i*_ = (*g*_*i*_, *b*) for some *b* ∈ *V* . It remains to show that *b* is uniquely determined. Let *N* (*g*_*i*_) denote the set of neighbors of *g*_*i*_. By Proposition 1, for exactly one element *h*^*′*^ ∈ *N* (*g*_*i*_), *w*^(*i−*1)^ is nonvanishing on 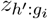; denote this element by *h*_*i*_. We claim *b* = *h*_*i*_. Suppose for contradiction that *b* = *h*^*′*^ ∈ *N* (*g*_*i*_) *\ {h*_*i*_*}*. Since *w*^(*i−*1)^ is nonvanishing on 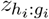, we have that *w*^(*i*)^ is nonvanishing on both 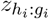 and 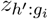. Therefore *w*^(*i*)^ is nonvanishing on 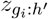, as 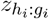 is a subtree of 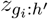. Therefore by Lemma 1, for some *j > i*, we have either *p*_*j*_ = (*g*_*i*_, *h*^*′*^) or *p*_*j*_ = (*h*^*′*^, *g*_*i*_), contradicting the bijection between edges and mutations. We conclude that 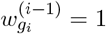 implies *b* = *h*_i_, so we have deterministically that *χ* = 0, *p* = (*g, h*), and

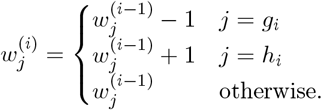

This corollary gives us a simple way of checking whether an evolutionary path *g* is valid or invalid: construct *w*^(*i*)^, *χ*_*i*_, and *p*_*i*_ for *i* = 1, 2, …, *m* + *n* 1 deterministically as in Corollary 1, and check at each iteration whether the axioms of a valid sequence of coalescences and mutations hold. If at any point they do not, or if Proposition 1 is ever violated, we may immediately declare *u* invalid. See Algorithm 1, which is essentially an equivalent formulation of the proof of Corollary 1, for details. As stated, this validity checking algorithm is polynomial in runtime: for each of the *m* iterations in which a mutation occurs, we must search all *m* nodes in the tree to determine the subtrees on which *w*^(*i−*1)^ vanishes. In practice, this complexity can be improved when *u* is a perturbation of an existing evolutionary path we already know to be valid. Specifically, by designing our MCMC sampler of evolutionary paths *u* in such a way that stores the unique (*χ, p*) corresponding to *u* at each iteration, as well as the functions *w*_0_, …, *w*_*m*+*n−*1_ corresponding to (*χ, p*), it is possible to verify in *O*(1) runtime whether swapping two consecutive elements of *u* preserves its validity. We omit the details here, but they amount to determining which neighboring coalescences and mutations are legal to swap based on the number of remaining lineages of each genotype they leave after having been performed.

In summary, we may restrict our MCMC algorithm to the space of valid evolutionary paths *u* by only allowing MCMC moves that preserve the validity of *u*. Using our adjacent transposition Metropolis-Hastings move, this validity check may be executed in *O*(1), allowing our sampler to achieve favorable statistical efficiency.

#### Algorithm 1

Checking the Validity of an Evolutionary Path

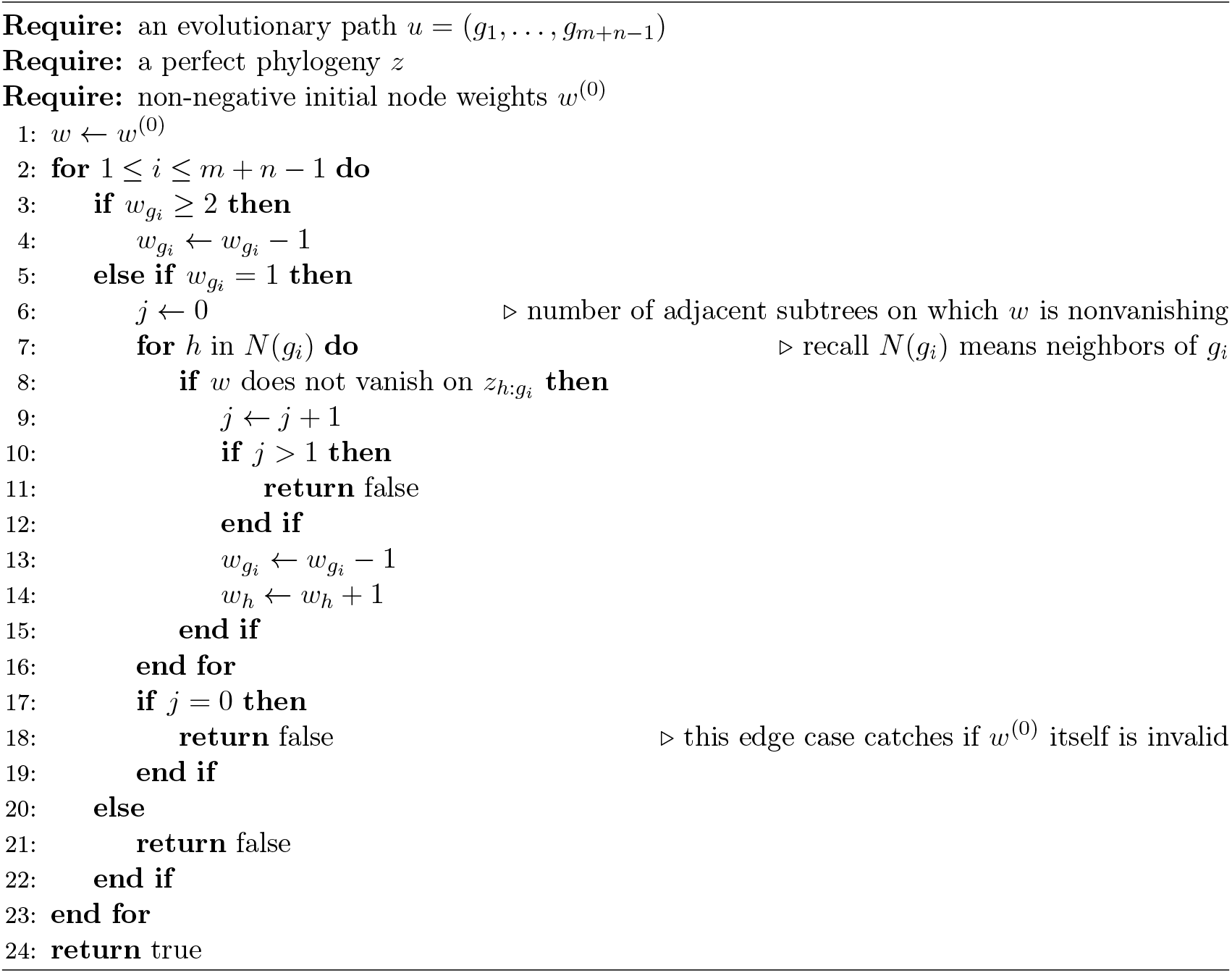

## B Proof of MCMC Irreducibility

Here, we prove that the adjacent transposition Markov chain proposed in Section 3.2 is irreducible, meaning that it explores the space of all valid evolutionary paths with positive probability. Recall that the chain is constructed by selecting an index *i* ∈ *{*1, …, *m* + *n −* 2*}* uniformly at random, and swapping *g*_*i*_ with *g*_*i*+1_ in an evolutionary path *u* = (*g*_1_, …, *g*_*m*+*n−*1_). To show irreducibility, we construct a specific evolutionary path *f*, and show that any evolutionary path *g* can be reached from *f* by a sequence of MCMC transposition moves.

We first define our initial valid evolutionary path *f* that is compatible with *z*. Consider a post-order traversal of nodes of *z* given by *h* = (*h*_1_, …, *h*_*m*+1_), i.e. an enumeration *h* of *V* such that for all integers 1 ≤ *i* ≤ *j* ≤ *k* ≤ *m* + 1, we have *d*(*h*_*j*_, *h*_*k*_) ≤ *d*(*h*_*i*_, *h*_*k*_), with *d* as defined in Appendix A. If the optional root *R* is specified, we further require *h*_*m*+1_ = *R*; if not, *h*_*m*+1_ may take on any value. A post-order traversal may be obtained by running depth-first search on *z* and then reversing the order in which nodes are discovered. If the optional root is specified, the depth-first search must start at *R*; otherwise, it may start at any node. We construct *f* = (*f*_1_, *f*_2_, …, *f*_*m*+*n−*1_) iteratively as follows: first, let *w*^(0)^ be as before, i.e. *w*^(0)^ is a tuple of *m* + 1 elements whose *j*th element 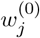 equals the number of sampled tips of genotype *j*. Set

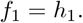

Now, iteratively for *i* ≤ *k* ≤ *m* + 1, suppose *f*_*i*_ = *h*_*k*_. Define *w*^(*i*)^ as in Corollary 1, i.e. *w*^(*i*)^ maps each genotype to its number of extant lineages after the *i*th coalescence or mutation in *f*_*i*_ in *f* . (See Appendix A, immediately prior to Definition 1 for the construction of *w*^(*i*)^.) We then set

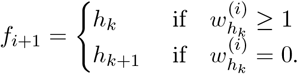

### Algorithm 2

Constructing a Valid Evolutionary Path

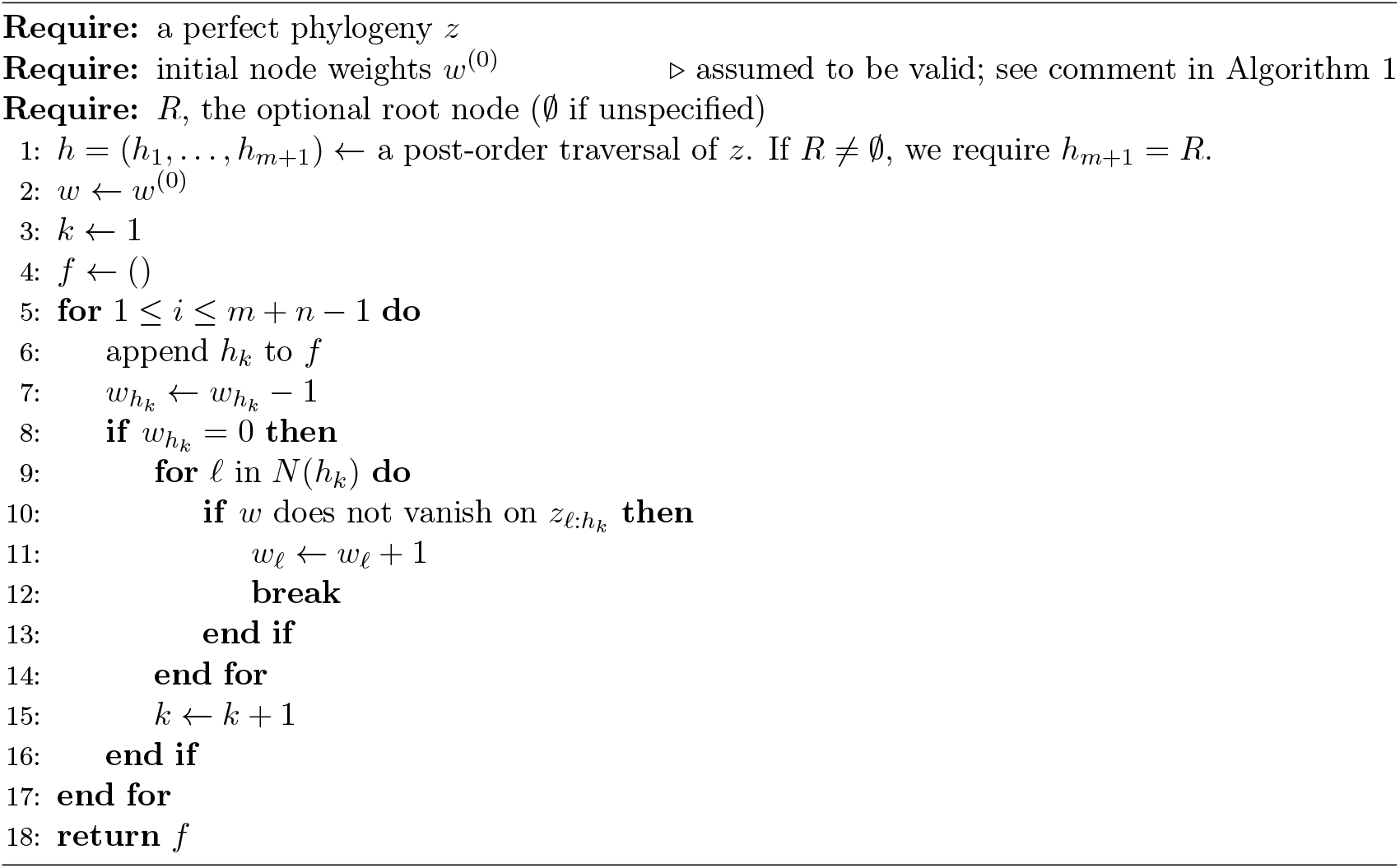

See Algorithm 2 for an equivalent formulation of this construction. Intuitively, the idea behind constructing *f* is that we start by considering a genotype observed at the tips, and apply coalescences within this genotype until there is only one left. We then apply a mutation to replace the one remaining lineage with a lineage of its ancestral genotype. We now move onto another genotype, none of whose descendant genotypes have any extant lineages, and apply the same procedure. We repeat this procedure until only one genotype with one extant lineage remains. If we follow the post-order traversal when choosing genotypes, it will always be true that each genotype has no descendant genotypes with extant lineages when the procedure is applied.

To see this formally, we claim that:

### Lemma 2

*f* is well-defined and is a valid evolutionary path.

*Proof*. We claim that *f* can be converted unambiguously to a sequence of coalescences and mutations using the algorithm established in Corollary 1. There are two components to proving this claim: first, that the two genotypes involved in each mutation event may be determined unambiguously, and second, that *h*_*k*+1_ in the defition of *f*_*i*+1_ is always well-defined, i.e. *k* + 1 never exceeds *m* + 1 (otherwise *f* itself is ill-defined).

We start with the first part of the claim. Consider *i* such that 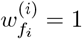 and *i < m* + 1, which means that event *i* + 1 must be a mutation in genotype *f*_*i*_ because there is exactly one extant lineage of that genotype after the *i*th event. We then have *f*_*i*+1_ = *f*_*i*_ by construction. Since *h* is a post-order traversal, *w*^(*i*)^ vanishes on all subtrees adjacent to *f*_*i*_ except for the one including *R*. Therefore, the two lineages involved in the mutation event indexed *i* + 1 are uniquely determined: one is *f*_*i*+1_, and the other is the unique neighbor of *f*_*i*+1_ located on the shortest path from *f*_*i*+1_ to *R*.

For the second part of the claim, we observe that *h*_*k*_ appears in *f* exactly 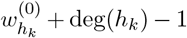 times, where deg(*h*_*k*_) is the degree of *h*_*k*_ (number of neighbors). To see this, first consider the case where *k*≠ *m* + 1. There are 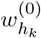 extant lineages of genotype *h*_*k*_ originating from the sampled leaves, as well as an additional deg(*h*_*k*_) *−* 1 extant lineages originating from the deg(*h*_*k*_) *−* 1 neighboring nodes of *h*_*k*_ visited earlier in the post-order traversal. These lineages require 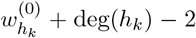 coalescences to be made into one extant lineage. This one extant lineage must then mutate into some neighboring genotype on the augmented perfect phylogeny, increasing the number of entries of *f* equal to *h*_*k*_ by 1. Therefore, 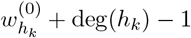 entries of *f* must equal *h*_*k*_.

If *k* = *m* + 1, once again, there are 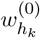 extant lineages of genotype *h*_*k*_ originating from the sampled leaves. Unlike before, there are deg(*h*_*k*_) extant lineages originating from the deg(*h*_*k*_) neighboring nodes of *h*_*k*_ visited earlier in the post-order traversal. In order for these 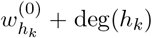 lineages to coalesce into one, 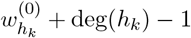 coalescent events are required. As no mutation is required for the final genotype in the post order traversal, we again find that *h*_*k*_ appears in *f* a total of 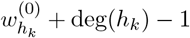 times.

Using this calculation, we have

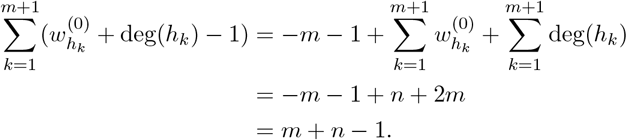

This shows the construvtion

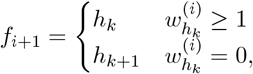

is well-defined, i.e. we never require the value of *h*_*k*+1_ for *k* + 1 *> m* + 1, because the *m* + *n* − 1 entries in *f* depend exactly on the values *h*_1_, …, *h*_*m*+1_. Now that we know *f* and its corresponding sequence of coalescences and mutations (*χ, p*) are well-defined, it follows by construction of (*χ, p*) as in Corollary 1 that the conditions of a valid sequence of coalescences and mutations are satisfied.

**Remark 1**. In the proof of the lemma, we showed that for all *k*, exactly 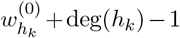 elements of *f* equal *h*_*k*_. The same reasoning can be used to show that for any *v* ∈ *V* and any evolutionary path *u, v* appears in *u* exactly 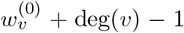 times. This observation confirms we need not sample evolutionary paths where the counts of entries of each genotype differ from *f*, as they are automatically invalid. Equivalently, the state space of valid evolutionary paths is in fact a subset of all permutations of *f* .

Finally, we claim that *f* can be achieved from any valid starting configuration of the Markov chain:

### Lemma 3

Let *u* = (*g*_1_, …, *g*_*m*+*n−*1_) be any valid evolutionary path. Then there exists a sequence of valid evolutionary paths *u* = *g*^(0)^, *g*^(1)^, …, *g*^(*p−*1)^, *g*^(*p*)^ = *f* such that *g*^(*i*+1)^ is identical to *g*^(*i*)^, except that for some *j*, the *j*th and (*j* + 1)st components of *g*^(*i*)^ appear swapped in *g*^(*i*+1)^.

*Proof*. Start with *u* = *g*^(0)^ and perform neighbor swaps (adjacent transposition) until all 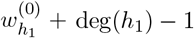 entries equal to *h*_1_ occupy the first 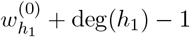 components of the evolutionary path. When performing the swaps, ensure that no swap increases the index of any entry of the evolutionary path equal to *h*_1_. This condition guarantees that the evolutionary path produced after each swap is valid, though there are also other possible ways to achieve the desired configuration. Then repeat for *h*_2_, performing swaps as necessary until the next 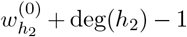 components of the evolutionary path (after those equal to *h*_1_) are equal to *h*_2_. Again, ensure that no swap increases the index of any entry of the evolutionary path equal to *h*_2_. Repeat again for *h*_3_, *h*_4_, …, *h*_*m*+1_ to achieve *f* . It is readily verified that every swap in this procedure maintains the validity of the evolutionary path.

### Proposition 2

The adjacent transposition Markov chain for exploring the space of evolutionary paths is irreducible.

*Proof*. Every adjacent transposition is proposed with positive probability, and every adjacent transposition that proposes a valid evolutionary path is accepted with positive probability under the Metropolis-Hastings criterion. By Lemmas 2 and 3, a finite sequence of adjacent transpositions can convert any valid evolutionary path into any other valid evolutionary path.

## C Laplace Approximation for Effective Population Size Inference

In this appendix, we derive a Multivariate Normal approximation to the density

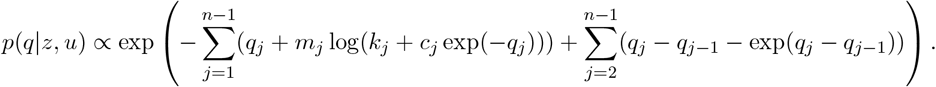

We first compute the mode of *p*(*q*|*z, u*) using numerical methods. Define

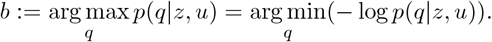

This optimization problem can be solved efficiently by Newton’s Method, as both the gradient and Hessian have analytic forms and the Hessian is tridiagonal. The gradient is given by

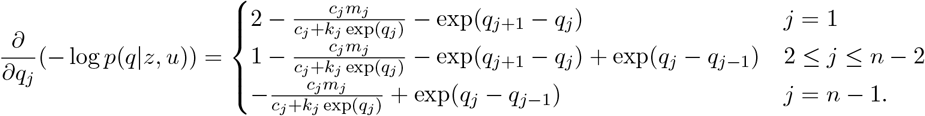

This Hessian has diagonal elements given by

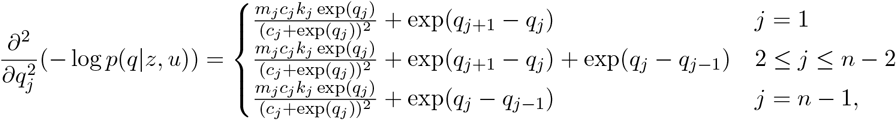

and off-diagonal elements given by

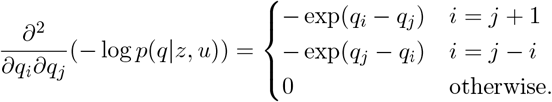

Thus, to propose a new value of *q* we first compute *b*. We then invoke the fact that if *p*(*q*|*z, u*) were, in fact, a multivariate normal density, its precision matrix would be given by the Hessian ∇^2^(™log *p*(*q*|*z, u*)). While this Hessian depends on *q*, it can be approximated to zeroth-order accuracy by setting *q* equal to its mode *b*, giving us a sampling density of

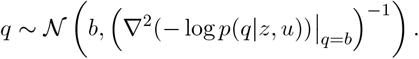

Since the above density and *p*(*q*|*z, u*) have the same Hessian at *q* = *b*, we have thus derived an approximate sampler that is second-order accurate about *q* = *b*. Moreover, *b* is a natural choice of location for which the approximation is most accurate, as it is in a region of high probability mass for *p*(*q*|*z, u*).

Thanks to the tridiagonal form of the Hessian, the entire proposal for *q* can be completed in *O*(*n*) operations, assuming the numerical maximization procedure of *p*(*q*|*z, u*) runs in *O*(1) iterations. This allows for a highly-efficient proposal effective population size trajectory.

## D Sequential Monte Carlo Tree Inference

Here we describe a sequential Monte Carlo (SMC) scheme for sampling trees in the form of evolutionary paths *u* conditional on the effective population size *N* . Recall that the density we wish to sample is given by:

**Figure S1:**
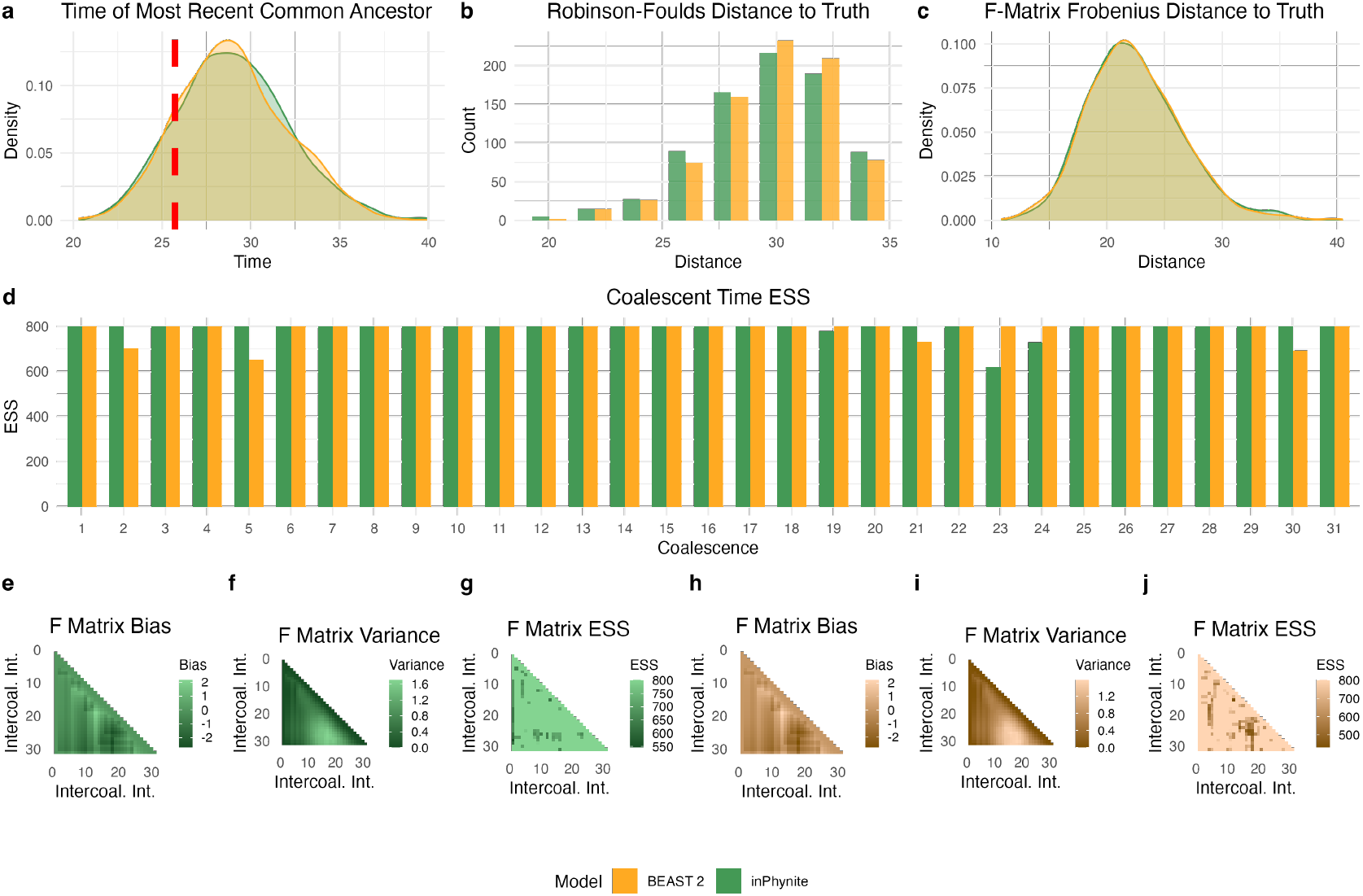
Performance comparison of `inPhynite` (top) to `BEAST 2` (bottom) for 32 taxa and effective population size *N*_*j*_ = 16 for each intercoalescent interval *j*. See caption of Figure 3 for more details.

**Figure S2:**
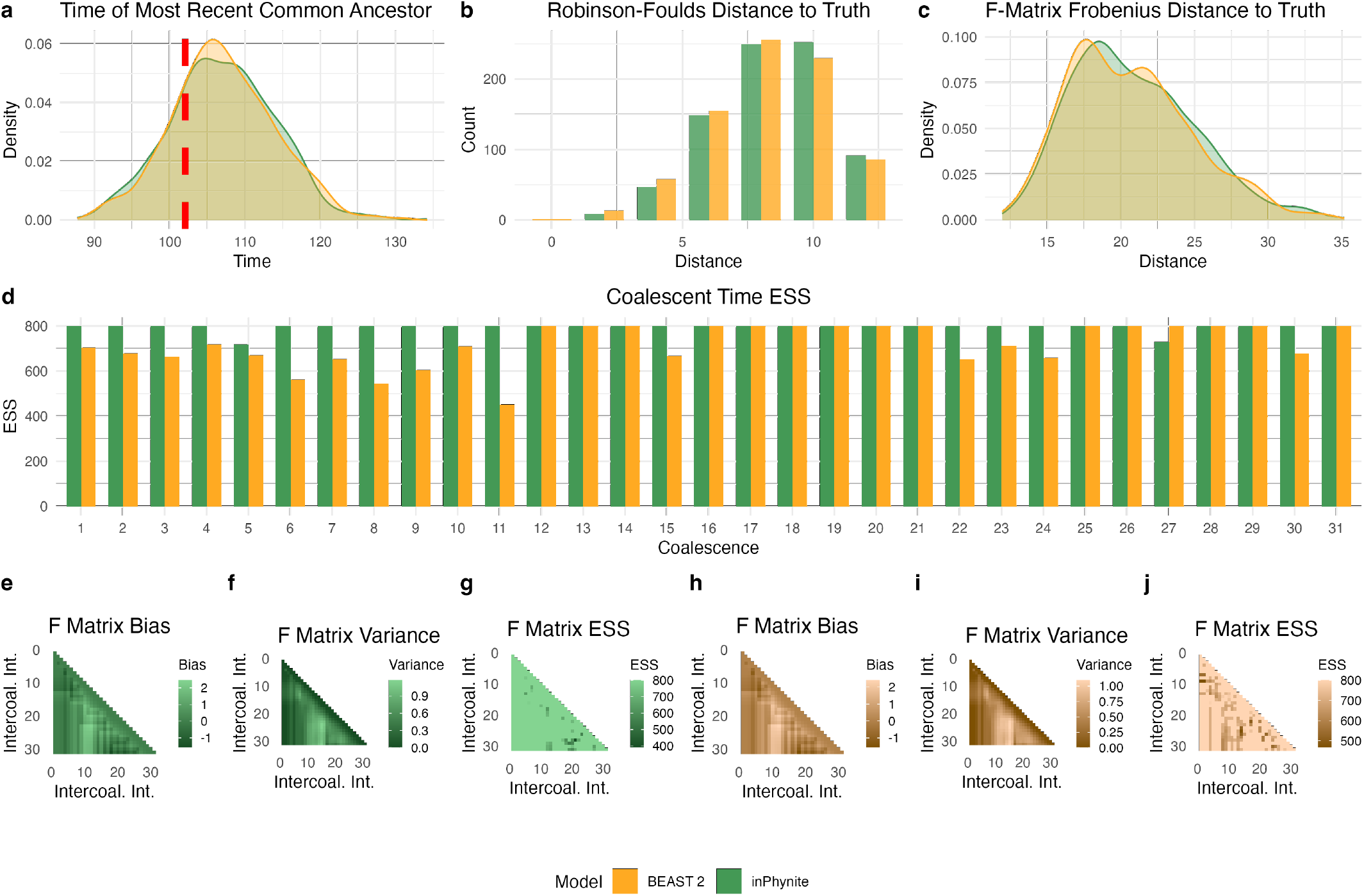
Performance comparison of `inPhynite` (top) to `BEAST 2` (bottom) for 32 taxa and effective population size *N*_*j*_ = 64 for each intercoalescent interval *j*. See caption of Figure 3 for more details.

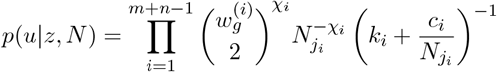

This is the probability density of a Markov process whose state space is the number of extant lineages of each genotype. Like in Appendix A, let 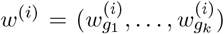 where *g*, …, *g* enumerates all genotypes in an augmented perfect phylogeny. To define the transition probabilities, recall from Appendix A that there are only two types of transitions with nonzero probability, and their density is given by

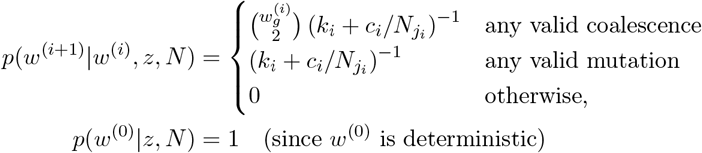

To see that *w*^(*i*)^ indeed captures all necessary information about the state space, we note that

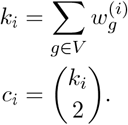

As the matrix *W* = (*w*^0^, …, *w*^*m*+*n−*1^) encodes identical information to an evolutionary path *u*, it is clear from the above that

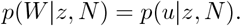

In addition to a Markovian decomposition of the target density, the other required component of Sequential Monte Carlo is a proposal distribution that is straightforward to sample. We denote this proposal density *m*(*w*^(*i*+1)^ | *w*^(*i*)^, *z, N*). We describe the proposal distribution in words; we then provide the mathematical form. For the coalescence or mutation to take place at iteration *i* + 1, we propose:

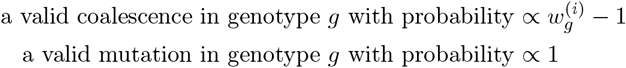

The transition kernel *m* may thus be written as

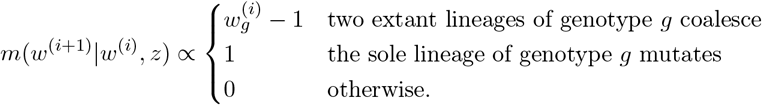

It is straightforward to calculate the normalizing constant of this proposal distribution by summing over genotypes *j* in which a coalescence or mutation event occurs. The calculation is not shown here.

The exact version of SMC we implemented is known as adaptive particle filtering (see Chopin and Papaspiliopoulos (2020), Algorithm 10.3), in combination with forward-filtering-backward-sampling (Chopin and Papaspiliopoulos (2020), Algorithm 12.2). To extend this method to a sampler for both trees and effective population size trajectories, we implemented particle Gibbs (Chopin and Papaspiliopoulos (2020), Algorithm 16.7). All code was implemented in C++ using the `OpenMP` library for CPU parallelization. With multithreading across up to 32 CPU cores, we were still unable to match the statistical efficiency of our Metropolis-Hastings sampler. As we observed in practice on the Yoruba dataset, after each particle Gibbs iteration, the value of *w*^(*i*)^ remained constant for approximately the first 80% of indices *i*, with only about the last 20% ever changing. This is a sign of severe particle degeneracy, an issue that could in theory be resolved with higher volumes of particles but would incur too great of a computational cost to be feasible.

## E Additional inPhynite–BEAST 2 Comparisons for Fixed Effective Population Size

Here, we compare the inferences of `inPhynite` and `BEAST 2` with the effective population size fixed at its true value. We show results on simulated datasets with 32 taxa and effective population sizes of 16 and 64 (assuming a mutation rate *µ* = 1). Like in Figure 3, posterior distributions agreed between the two methods, and both achieved effective sample sizes indicative of good convergence.

## F Additional inPhynite–BEAST 2 Comparisons for Variable Effective Population Size

Here, we compare the inferences of `inPhynite` and `BEAST 2` with inferred effective population sizes, using simulated data generated under the “constant” and “exponential decay” regimes outlined in Section 4.2. Like in Figure 4, posterior distributions agreed between the two methods, and both achieved effective sample sizes indicative of good convergence. These results were achieved in less than one minute for `inPhynite`, but took over 20 minutes for `BEAST 2`.

**Figure S3:**
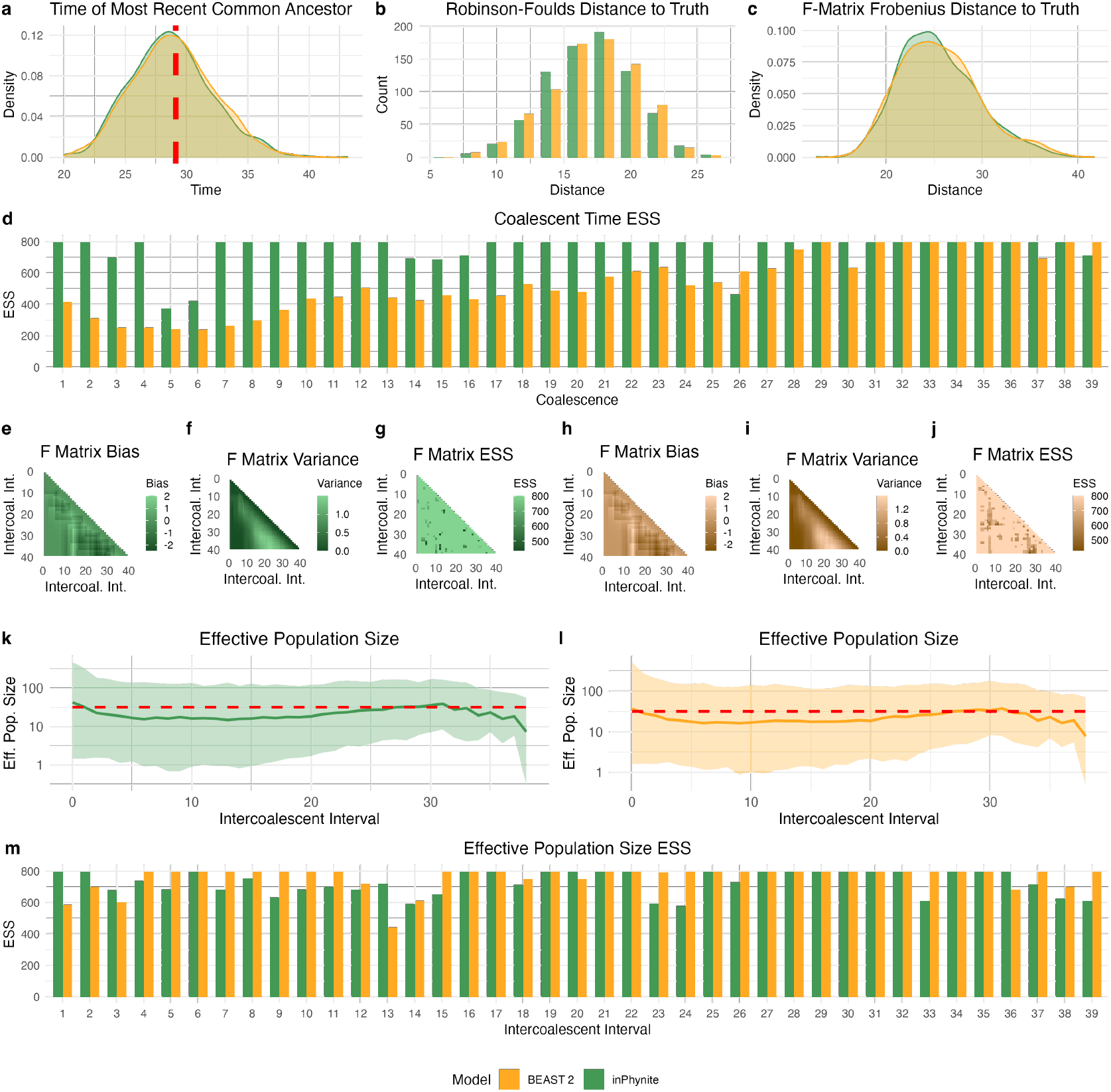
`inPhynite` and `BEAST 2` inferences and performance for a constant effective population size trajectory. See caption of Figure 4 for more details.

**Figure S4:**
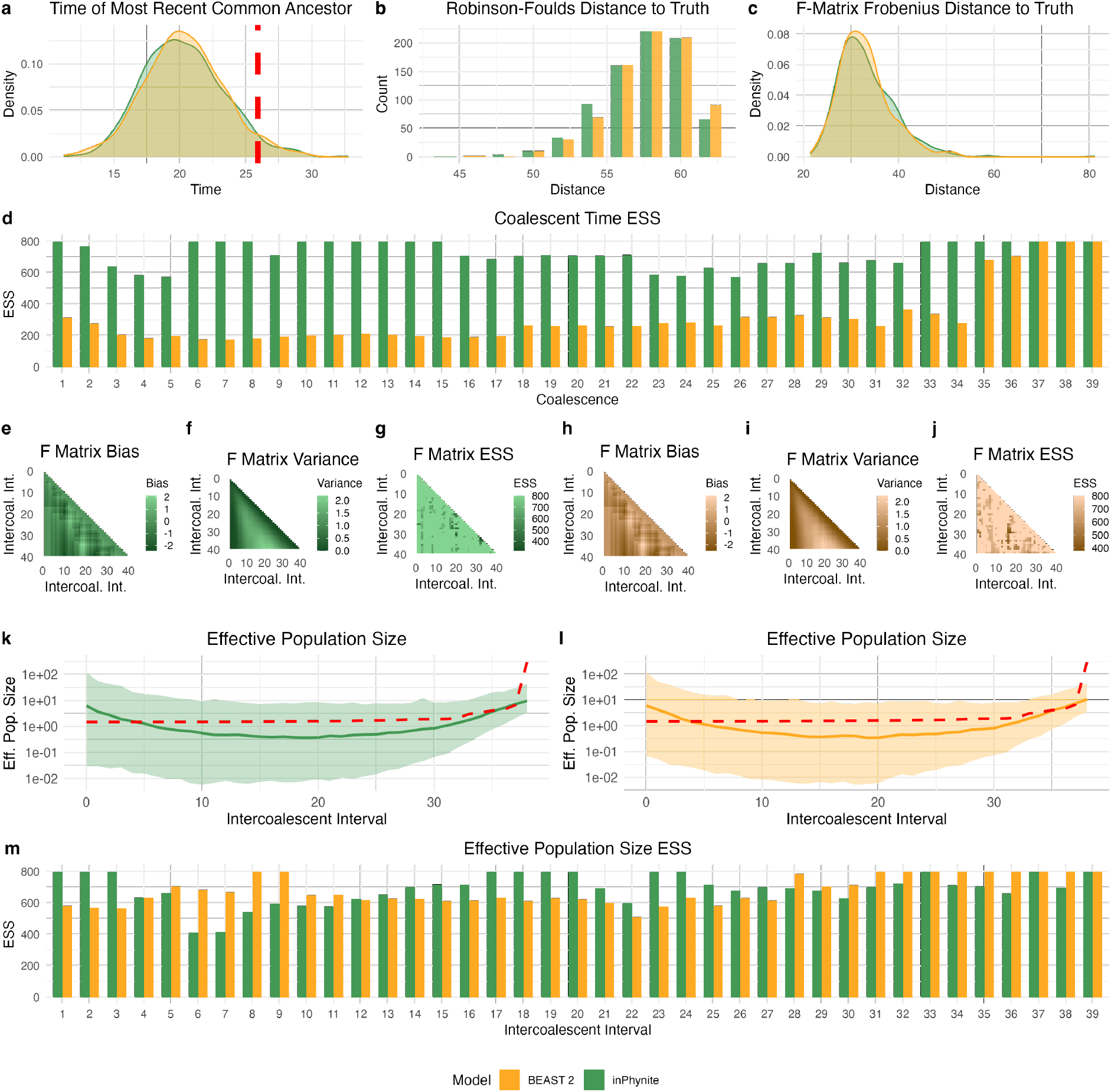
`inPhynite` and `BEAST 2` inferences and performance for an exponentially decaying effective population size trajectory. See caption of Figure 4 for more details.

